# Excitatory GABAergic signalling is associated with acquired benzodiazepine resistance in status epilepticus

**DOI:** 10.1101/478594

**Authors:** Richard J. Burman, Joshua S. Selfe, John Hamin Lee, Maurits van den Burg, Alexandru Calin, Neela K. Codadu, Rebecca Wright, Sarah E. Newey, R. Ryley Parrish, Arieh A. Katz, Joanne M. Wilmshurst, Colin J. Akerman, Andrew J. Trevelyan, Joseph V. Raimondo

## Abstract

Status epilepticus (SE) is defined as a state of unrelenting seizure activity. Generalised convulsive SE is associated with a rapidly rising mortality rate, and thus constitutes a medical emergency. Benzodiazepines, which act as positive modulators of chloride (Cl^-^) permeable GABA_A_ receptors, are indicated as first-line treatment, but this is ineffective in many cases. We found that 48% of children presenting with SE were unresponsive to benzodiazepine treatment, and critically, that the duration of SE at the time of treatment is an important predictor of non-responsiveness. We therefore investigated the cellular mechanisms that underlie acquired benzodiazepine resistance, using rodent organotypic and acute brain slices. Removing Mg^2+^ ions leads to an evolving pattern of epileptiform activity, and eventually to a persistent state of repetitive discharges that strongly resembles clinical EEG recordings of SE. We found that diazepam loses its antiseizure efficacy and conversely exacerbates epileptiform activity during this stage of SE-like activity. Interestingly, a low concentration of the barbiturate phenobarbital had a similar exacerbating effect on SE-like activity, whilst a high concentration of phenobarbital was effective at reducing or preventing epileptiform discharges. We then show that the persistent SE-like activity is associated with a reduction in GABA_A_ receptor conductance and Cl^-^ extrusion capability. We explored the effect on intraneuronal Cl^-^ using both gramicidin, perforated-patch clamp recordings and Cl^-^ imaging. This showed that during SE-like activity, reduced Cl^-^ extrusion capacity was further exacerbated by activity-dependent Cl^-^ loading, resulting in a persistently high intraneuronal Cl^-^. Consistent with these results, we found that optogenetic stimulation of GABAergic interneurons in the SE-like state, actually enhanced epileptiform activity in a GABA_A_R dependent manner. Together our findings describe a novel potential mechanism underlying benzodiazepine-resistant SE, with relevance to how this life-threatening condition should be managed in the clinic.

## Introduction

The majority of all spontaneously occurring seizures terminate within a few seconds to minutes, and without medical intervention. When seizures fail to stop naturally, this is referred to as status epilepticus (SE). This represents a neurological emergency and requires immediate therapeutic intervention. Convulsive SE occurs more frequently in children, and if not managed effectively, is associated with significant morbidity and even mortality (Boggs, 2004). Current first-line treatment for SE recommends the use of benzodiazepines (Glauser *et al*., 2016). These drugs work by positively modulating Cl^-^-permeable ionotropic GABA_A_ receptors (GABA_A_R), which underlie the majority of fast inhibitory neurotransmission within the brain. The intent is to boost inhibitory signalling in an attempt to terminate SE. Unfortunately, benzodiazepine treatment fails to terminate seizures in a large fraction of patients, which underscores the inadequacy of our current first-line therapeutic strategy for treating this condition (Appleton *et al*., 2000; Mayer *et al*., 2002; Chin *et al*., 2008).

Current thinking in the field is that benzodiazepine resistance in SE is largely due to impaired GABA_A_R trafficking (Goodkin and Kapur, 2009). Both *in vitro* and *in vivo* animal models have shown that extended seizure activity is correlated with internalisation of GABA_A_Rs in the hippocampus (Kapur and Coulter, 1995; Goodkin *et al*., 2005; Naylor *et al*., 2005). In addition, SE has been shown to be associated with a reduction in surface expression of GABA_A_R subunits (γ2), which are necessary for benzodiazepine binding (Goodkin *et al*., 2008). This line of reasoning suggests that SE-induced changes to the GABA_A_R impairs the ability of benzodiazepines to enhance the GABA_A_R conductance, thereby resulting in treatment failure.

It is well accepted that the intracellular concentration of Cl^-^, and therefore the reversal potential for GABA_A_Rs (E_GABA_) can change over multiple time scales (Wright *et al*., 2011; Ellender *et al*., 2014; Sato *et al*., 2017). Long-term changes in the expression of Cl^-^ transporter proteins modifies steady-state E_GABA_ over development and in multiple disease states including epilepsy (Moore *et al*., 2017). In addition to these well-described long-term changes, short-term (seconds to minutes) changes in E_GABA_ can occur following intense GABA_A_ activation that causes Cl^-^ influx, which can overwhelm Cl^-^ extrusion mechanisms (Alger and Nicoll, 1979; Kaila *et al*., 1989; Staley *et al*., 1995; Wright *et al*., 2011). Furthermore, significant Cl^-^ accumulation and a temporary excitatory shift in GABAergic signalling has been shown to accompany single seizure-like events in both *in vitro* (Lamsa and Kaila, 1997; Isomura *et al*., 2003; Fujiwara-Tsukamoto *et al*., 2010; Ilie *et al*., 2012; Ellender *et al*., 2014) and *in vivo* (Sato *et al*., 2017) models. The breakdown of the Cl^-^ gradient serves to explain the surprising excitatory effects of GABAergic interneuronal subtypes recently observed during *in vitro* and *in vivo* seizure events (Ellender *et al*., 2014; Khoshkhoo *et al*., 2017; Magloire *et al*., 2019). This suggests that in circuits where seizures have caused sufficient Cl^-^ loading, GABA_A_R mediated synaptic signalling can serve to exacerbate rather than control hyperexcitablity, which has direct relevance for the inhibitory efficacy of benzodiazepines. Deeb *et al*. (2013) have previously demonstrated that the benzodiazepine diazepam has reduced inhibitory capability under these conditions of activity-driven Cl^-^ accumulation. However, it is currently unknown as to whether Cl^-^ dynamics and seizure-associated shifts in GABAergic signalling are involved in the development of benzodiazepine resistance in SE.

In this study we combine clinical and experimental data to explore the phenomenon of benzodiazepine-resistant SE. First we document the incidence of benzodiazepine resistance in a South African cohort of paediatric patients in SE and provide clinical evidence that seizure duration prior to treatment is a useful predictor of benzodiazepine resistance. As SE is most often caused by acute brain insults resulting in sustained seizure-activity, we used the acute *in vitro* 0 Mg^2+^ model of SE (Dreier *et al*., 1998) in both organotypic and acute brain slices to explore the cellular mechanisms underlying acquired benzodiazepine resistance. We demonstrate that the benzodiazepine diazepam loses its antiseizure efficacy and can actually enhance epileptiform discharges during SE-like activity. Similarly, we find that a low concentration of the barbiturate phenobarbital also exacerbates SE-like activity, whilst a high concentration of phenobarbital maintains an antiseizure effect. Utilising gramicidin perforated patch-clamp recordings, Cl^-^ imaging and optogenetic control of GABAergic interneurons, we characterise changes in intracellular Cl^-^ and GABAergic signalling during the development of SE. We find that although GABA mediated conductances are reduced in early SE, pharmacoresistance is associated with deficits in Cl^-^ extrusion capability and profound activity-dependent Cl^-^ accumulation which results in excitatory interneuronal signalling via GABA_A_Rs.

## Materials and methods

### Clinical data

Clinical data was obtained from paediatric patients presenting with convulsive status epilepticus (CSE) to the Red Cross War Memorial Children’s Hospital (RCWMCH) from 2015 to 2018 as part of a clinical trial comparing second-line therapy for paediatric status epilepticus in a resource-limited setting. This study was approved by the University of Cape Town Human Ethics Commiittee (HREC 297/2005) and the study protocol was registered on the ClinicalTrials.gov registry (NCT03650270). The findings of this study have been published (Burman *et al*., 2019). CSE was defined as any seizure that lasts longer than five minutes, or multiple discrete seizures between which there is no extended period of recovery between events (Trinka *et al*., 2015). The onset of CSE was taken as the time when the child caregiver or healthcare professional first documented clinical signs of a convulsive seizure. Upon admission, all patients were treated with benzodiazepines. If CSE continued after two doses of benzodiazepines, patients were then given second-line therapy. A successful response to treatment was defined as a termination of signs of convulsive seizure activity. If a child did not respond to first-line treatment with benzodiazepines, they then received one of two second lines regimens. Study data was collected using a custom-made REDCap database (hosted by the University of Cape Town’s eResearch Centre).

### Brain slice preparation

Slices were prepared from GAD2-cre-tdTomato mice (C57BL/6 background, JAX lab) or wistar rats. The GAD2-cre-tdTomato strain which resulted in cre-recombinase and tdTomato expression in all GABAergic interneurons (Taniguchi *et al*., 2011). The use of animals was approved by the University of Cape Town Animal Ethics Committee (mouse) or in accordance with regulations from the United Kingdom Home Office Animals (Scientific Procedures) Act (rat). Organotypic brain slices were prepared using 7 day old animals and followed the protocol originally described by Stoppini *et al*. (1991) (for details see Raimondo *et al*. (2016)). Recordings were performed 6-14 days post culture which is equivalent to P13 to P21. This and previous work has shown that pyramidal neurons in the organotypic hippocampal brain slice have mature and stable Cl^-^ homeostasis mechanisms at this point, as evidenced by their hyperpolarizing E_GABA_ (Ilie *et al*., 2012; Raimondo *et al*., 2012). For experiments using optogenetics, after 1 day in culture, slices prepared from GAD2-cre-tdTomato mice were transduced with adeno-associated vector serotype 1 (AAV1) containing a floxed-STOP channelrhodopsin (ChR2) linked to a yellow fluorescent protein (YFP), which resulted in selective expression of ChR2 in GABAergic interneurons (Royo *et al*., 2008). For Cl^-^ imaging experiments, neurons were biolistically transduced with ClopHensorN construct following the same procedure described by Raimondo *et al*. (2013). Acute brain slices were prepared from P14-P21 mice. Horizontal slices of the temporal lobe included the entorhinal cortex and hippocampal formation (as demonstrated in Mann *et al*. (2009) and shown in Supp. Fig. 2).

### Electrophysiology

Brain slices were transferred to a submerged recording chamber (whole-cell and perforated patch experiments) or an interface recording chamber (LFP experiments) where they were continuously superfused with standard artificial CSF (aCSF) bubbled with carbogen gas (95% O_2_: 5% CO_2_) using peristaltic pumps (Watson-Marlow). The standard aCSF was composed of (in mM): NaCl (120); KCl (3); MgCl2 (2); CaCl2 (2); NaH2PO4 (1.2); NaHCO3 (23); D-Glucose (11) with pH adjusted to be between 7.35-7.40 using 0.1 mM NaOH. For patch-clamp experiments, neurons were visualized using a BX51WI upright microscope (Olympus) using 20x or 40x water-immersion objectives and targeted for recording. For whole-cell recordings, micropipettes were prepared from borosilicate glass capillaries (Warner Instruments) and filled with a low Cl^-^ internal solution composed of (in mM): Kgluconate (120); KCl (10); Na_2_ATP (4); NaGTP (0.3); Na_2_-phosphocreatinine (10) and HEPES (10). When recording GABAergic synaptic currents, pipettes were filled with a high Cl^-^ internal solution (Cl^-^ 141mM) composed of (in mM): KCl (135), NaCl (8.9) and HEPES (10). Gramicidin perforated patch recordings (Kyrozis and Reichling, 1995) were performed using glass pipettes containing the high Cl^-^ internal solution. Adequate perforation of the membrane was assessed by monitoring access resistance and was defined when access resistance was <90 MΩ. Patch-clamp recordings were made with Axopatch 200B amplifiers (Molecular Devices) and data acquired using WinWCP (University of Strathclyde). LFP recordings were performed using an AC-coupled amplifier (AMSystems). Data was acquired using the LabChart Pro (AD Instruments) with recordings processed using a 140Hz low-pass filter. SLEs were defined as events where significant deviations from the resting potential in LFP and patch-clamp recordings (>2 standard deviations) lasting at least 5 seconds. The LRD phase was defined as recurrent epileptiform discharges that persisted for at least 5 minutes. Slices that developed spontaneous SLEs prior to 0 Mg^2+^ exposure were excluded from analysis in order to ensure that prior epileptiform activity would not have altered the neuronal network (Kamphuis *et al*., 1991; Morimoto *et al*., 2004). The following drugs were used: diazepam; flumazenil; CGP-35348; kynurenic acid; tetrodotoxin, QX-314 (Tocris) and phenobarbital (Aspen Pharmacare).

### Confocal imaging

A confocal microscope (Zeiss) was used to visualise tdTomato and YFP-labelled ChR2 expression using 561 nm and 488 nm lasers. ClopHensorN (modified from the original ClopHensor by Arosio *et al*. (2010)) expressing neurons were excited using 458, 488 and 561 nm lasers and emission collected by photomultiplier tubes (PMTs): between 500 - 550 nm for EGFP and 635 - 700 nm for the tdTomato. Calibration of the reporter and Cl^-^ measurements were made as previously described Raimondo *et al*. (2013).

### Cell surface biotinylation and western blotting

Organotypic hippocampal slices were incubated for 3 hours at 28°C–30°C, in either control aCSF or aCSF with 0 Mg^2+^, while continuously bubbling with 95% O_2_-5% CO_2_. Biotinylation was performed to produce “total” and “surface” protein lysate samples as described in (Wright *et al*., 2017). Blots were incubated with rabbit anti-C-terminus KCC2 (1:500, Merck Millipore) followed by horseradish peroxidise (HRP)-conjugated goat anti-rabbit antibody (1:2000, Thermo Scientific). For each sample, the surface protein was normalized against the total protein, which was run in the adjacent lane. As the ratio of surface/total was calculated within each sample, this controlled for differences in overall protein levels across samples and variance associated with loading.

### Data analysis

Clinical data was analysed using SPSS Statistics (IBM) while experimental data analysis was performed using Matlab (MathWorks). Statistical measurements were performed using GraphPad Prism. Data reported as either median with interquartile range (IQR) or mean ± standard error of the mean unless otherwise stated.

### Data availability statement

Upon publication, all data presented in this manuscript will be made available on an open-access data repository.

## Results

### Benzodiazepine-resistance in status epilepticus is associated with enhanced morbidity and can be predicted by seizure duration prior to treatment

A total of 144 admissions of paedaitric CSE from 111 patients were observed at RCWMCH between 2015-2018 (median age at admission 28.11 months *IQR* 15.5 – 66.1). 52% of admissions responded to first-line benzodiazepine treatment, typically lorazepam, diazepam or midazolam (termed ‘benzodiazepine-sensitive’ or ‘bzpS’), while 48% did not (termed ‘benzodiazepine-resistant’ or ‘bzpR’) (Fig. 1A). We found no association between whether a patient had previous been admitted for seizures and benzodiazepine sensitivity, nor in whether they were previously diagnosed as having epilepsy (**Supp. Table 1**). For patients admitted multiple times, we found no difference in benzodiazepine sensitivity (**Supp. Table 2**) Benzodiazepine resistance was associated with enhanced morbidity as bzpR patients required longer care in hospital (Fig. 1B) and were more likely to need admission to the paediatric intensive care unit (PICU, Fig. 1C). Whilst acute causes were the most common precipitant of SE, we found no difference in the underlying aetiology between the groups (Fig. 1D). We found no significant difference between age at admission (**Supp. Table 3**). Furthermore, we found no association between CSE type, semiology or diagnosis of febrile status epilepticus and benzodiazepine sensitivity (**Supp. Table 3**).

**Figure 1:**
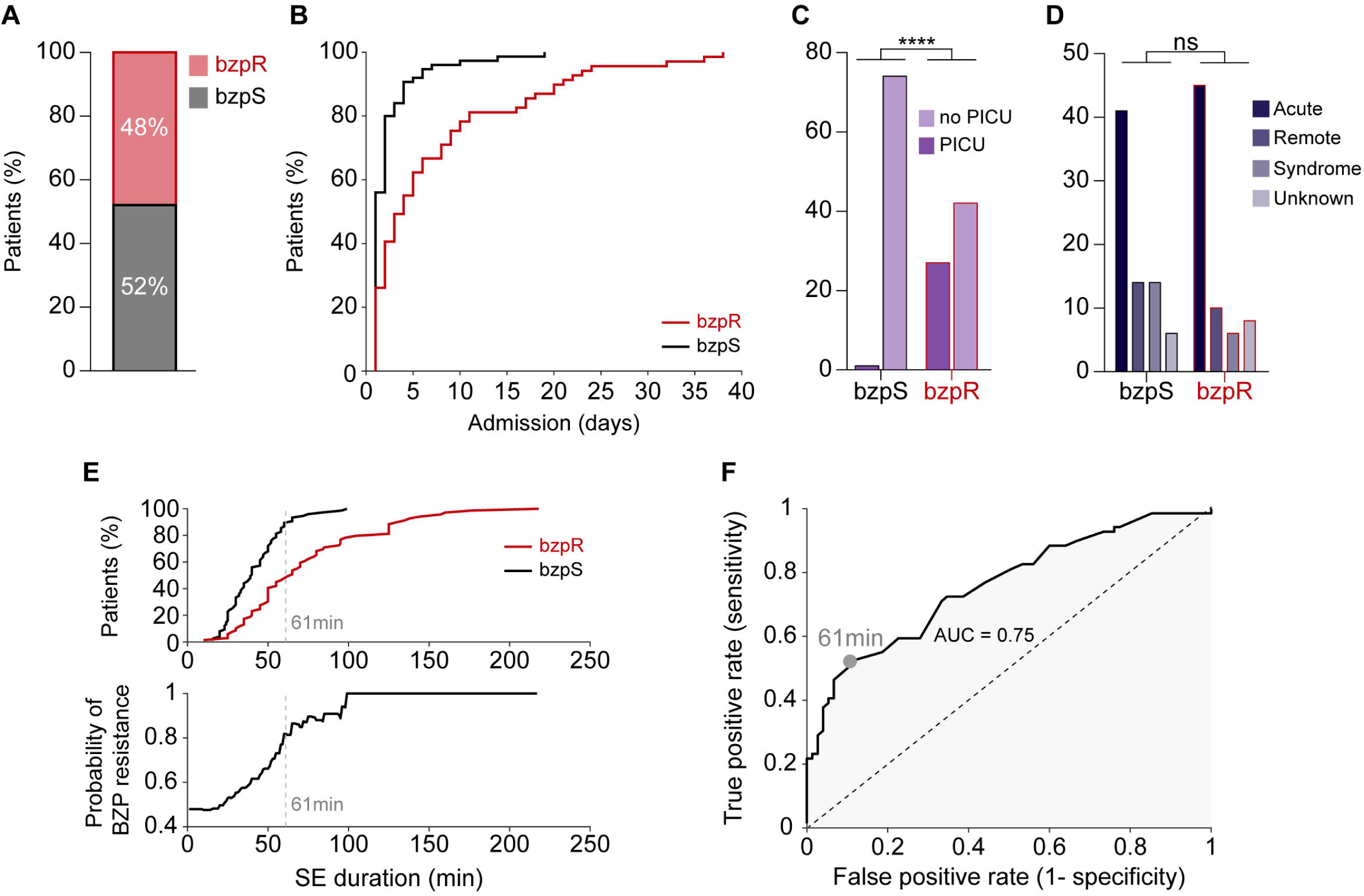
Resistance to first-line benzodiazepine treatment increases with the duration of status epilepticus and is associated with increased morbidity. **A**, Proportion of paediatric patients presenting with convulsive status epilepticus (CSE) resistant (bzpR, red) or sensitive (bzpS, black) to first-line treatment with benzodiazepines. **B**, Benzodiazepine resistance was associated with longer hospital stays (bzpS: median 4.00 *IQR* 1.0 – 2.0 days vs bzpR: median 4.0 days *IQR* 1.0 - 9.50 days, *p* < 0.0001, *Mann-Whitney U test*) and **C**, these patients were more likely to require admission to the paediatric intensive care unit (PICU, *OR* = 47.57, *95% CI:* 6.24 – 362.9, *p* < 0.0001, *Fisher-exact test,*). **D,** There, was no difference in underlying aetiology between the groups when dividing causes into four categories: ‘Acute’, acute illness; ‘Remote’, previous brain injury; ‘Syndrome’, established electroclinical syndrome; ‘Unknown’, no aetiology found during admission (*p* = 0.25, *Chi-Squared test*). **E**, Top, cumulative frequency plot of bzpR (red) and bzpS (black) patients as a function of seizure duration prior to treatment (bzpR: median 40.0 *IQR* 28.0 – 52.0 min vs bzpS: median 65.00 *IQR* 45.0 – 95.0 min, *p* < 0.0001, *Mann-Whitney U test*). Bottom, increased SE duration is associated with enhanced probability of benzodiazepine resistance. The optimum discrimination threshold for separating bzpR from bzpS patients calculated using ‘**F**’ is indicated by the grey dashed line. **F**, ROC curve, analysing how well the SE duration classifies the likelihood of benzodiazepine sensitivity or resistance. Top right corresponds to short SE durations; bottom left to long SE durations. The maximal difference between the true and false positive rates occurs when the classifier is set at 61 mins (grey circle). Error bars indicate mean ± SEM.

The median duration of SE prior to the time of initial treatment was 50.0min (*IQR* 33.0 – 69.5), but notably, longer SE durations were increasingly associated with benzodiazepine resistance (Fig. 1E). These data suggested that the duration of SE at the time of admission could be a useful clinical classifier, especially since benzodiazepine resistance is associated with enhanced morbidity. We addressed this question using receiver operating characteristic (ROC) curve analysis (Fig. 1F), by assessing the relative proportions of true positive rates (short duration SE are bzpS; long duration SE are bzpR) to false positive rates, as the classifier time changes from short (top right) to long (bottom left) durations. The area under curve (AUC) of 0.75 indicates that SE duration does indeed discriminate between these two patient populations, and importantly, it further indicates that the optimum discrimination threshold occurs at almost exactly 1 hour (maximal deviation from the diagonal was at 61 min; Fig. 1E, F). A contingency table (“confusion matrix”) using 1 hour as the cut off (see **Supp. Table 4**), shows that prior to this time, 67% of patients responded to benzodiazepine, but after that, only 18% did.

These clinical data suggest strongly that a critical factor is the continued seizure activity itself, and that the benzodiazepine resistance is an acquired, activity-dependent phenotype. There are striking parallels of pharmacoresistance, acquired over a similar time course, in acute rodent *in vitro* models, and so we used such a model to explore the underlying cellular mechanism.

### Early application of diazepam is antiseizure whilst late application enhances epileptiform activity in an in vitro model of status epilepticus

Our clinical data demonstrated a diverse set of causes for SE, and no particular set of causes of SE which enhanced the likelihood of benzodiazepine resistance. As acute brain insults (the most common cause in our clinical cohort) causing extended seizures lasting minutes to hours can result in benzodiazepine resistance, we used the well characterised *in vitro* 0 Mg^2+^ model of acute seizures and SE (Anderson *et al*., 1986; Mody *et al*., 1987; Gutiérrez and Heinemann, 1999; Albus *et al*., 2008). Here Mg^2+^ removal from the artificial cerebrospinal fluid (aCSF), results in initial interictal-like activity followed by the gradual development of seizure-like events, which mimic what is observed in temporal lobe seizures in humans (Anderson *et al*., 1986; Dreier *et al*., 1998). Importantly, after extended periods of Mg^2+^ withdrawal, distinct ictal events no longer occur and are replaced by persistent seizure-like activity in the form of recurrent epileptiform discharges (Anderson *et al*., 1986; Dreier *et al*., 1998), which strongly resemble clinical EEG recordings of convulsive SE (Fig. 2A). This late-stage activity is also referred to as the late recurrent discharge phase (LRD) and represents the best available *in vitro* model of SE (Zhang *et al*., 1995; Dreier *et al*., 1998). We utilised the 0 Mg^2+^ model of SE in two *in vitro* preparations; organotypic hippocampal slice cultures and acute slices of temporal cortex. Although organotypic slices were more excitable than the acute slices, with a greater propensity to generate SLEs and LRD (Supp. Fig. 2A-D), both preparations generated the well described transition from SLEs to SE-like LRD activity (Fig. 2A, Supp. Fig. 2G). We therefore sought to determine the effect of diazepam on the evolution of *in vitro* seizure-like activity using both these preparations.

**Figure 2:**
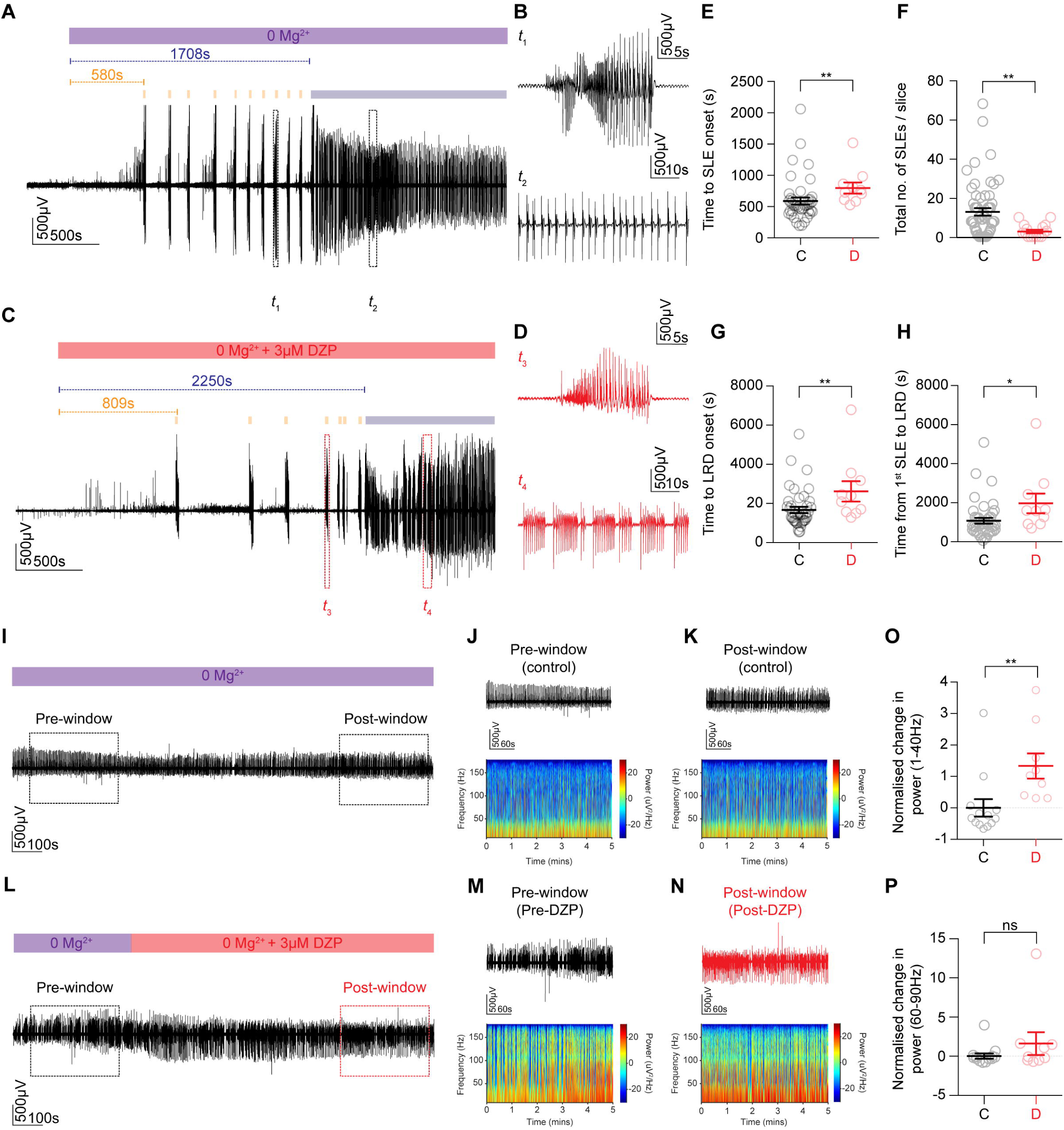
Early application of diazepam has an antiseizure effect while late application is ineffective and augments epileptiform bursting activity in an *in vitro* model of status epilepticus. **A**, The 0 Mg^2+^ chemoconvulsant model was used as an *in vitro* model of status epilepticus (SE) in organotypic hippocampal brain slices. LFP recording from CA1 demonstrating that upon Mg^2+^ withdrawal, activity progressed from single seizure-like events (SLEs, orange bars) to a phase where recurrent discharges occurred unabated, the late recurrent phase (LRD, blue bar). **B,** Window *t*_1_ from ‘A’ depicting a single SLE (top trace) and window *t*_2_ showing recurrent discharges or SE-like activity (bottom trace). **C**, Early application of diazepam (3 μM) introduced when 0 Mg^2+^ was washed in (red bar). **D**, Windows *t*_1_ and *t*_2_ depicting a SLE (top trace) and SE-like activity (bottom trace) in diazepam. **E**, Population data showing that early diazepam application delays the onset of SLEs (control, *n* = 41: median 510.0 *IQR* 412.6 – 616.2 s vs diazepam, *n* = 10: median 736.3 *IQR* 594.4 – 883.0 s, *p* = 0.003, *Mann-Whitney U test*) while decreasing the total number of SLEs (**F**, control: median 9.0 *IQR* 2.3 – 19.3 vs diazepam: median 2.0 *IQR* 0.0 – 5.8, *p* = 0.001, *Mann-Whitney U test*), retarding onset of LRD (**G**, control: median 1386 *IQR* 1087 – 1966 s vs diazepam: 2220 *IQR* 1522 – 3236 s, *p* = 0.01, *Mann-Whitney U test*) and extending the time from 1^st^ SLE to LRD (**H**, control: median 799.8 *IQR* 501.4 – 1228 s vs diazepam: median 1555sIQR 874.2 – 2529 s, *p* = 0.01, *Mann-Whitney U test*). **I,** LFP recording of the LRD phase in a control slice, with a 5min ‘Pre-window’ and ‘Post-window’ used for analysis (dashed rectangles). **J,** ‘Pre-window’ LFP trace (top) with its associated spectrogram (bottom). **K**, ‘Post-window’ as in ‘**J**’. **L,** LFP recording of LRD with diazepam application and accompanying ‘Pre-window’ (**M**) and ‘Post-window’(**N**) (dashed rectangles) used for analysis. **O,** Population data demonstrating raised normalised change in power of SE-like activity in the 1-40 Hz frequency range between diazepam and control slices (control, *n* = 13: median −0.3 *IQR* −0.5 - −0.01vs diazepam, *n* = 9: median 1.0 *IQR* 0.3 – 2.2, *p* = 0.001, *Mann-Whitney U-test*). **P,** No statistical difference in normalised change in power between groups was observed in the 60-90 Hz range (control: median −0.3 *IQR* −0.6 - −0.02 vs diazepam: median −0.18 *IQR* −0.6 – 1.5, *p* = 0.4, *Mann-Whitney U test*). ‘DZP’, diazepam. \**p* ≤ 0.05; \*\**p* ≤ 0.01; ‘ns’, not significant (*p* ≥ 0.05); error bars indicate mean ± SEM.

First, we showed that diazepam (3 μM) increases the decay time constant (tau) but not the amplitude of voltage-clamp recorded GABA_A_R synaptic currents (GSCs) elicited via optogenetic activation of GABAergic interneurons in organotypic brain slices (Supp. Fig. 1A-F). Furthermore, we then show that diazepam increases both the amplitude and decay time constant of GSCs elicited via electrical stimulation of afferent fibres (3 ms) in acute brain slices in the presence of the glutamate receptor blocker, kynurenic acid (2 μM) Supplementary Fig. 1G-J). In both these preparations the effects of diazepam could be reversed by application of its competitive antagonist, flumazenil (0.4 μM). These findings confirm that GABAergic signalling in both organotypic and acute brain slices is sensitive to diazepam during baseline activity.

We then monitored the evolution of 0 Mg^2+^ induced seizure activity using local field potential recordings on interface, whilst diazepam (3 μM), equivalent to a total clinical dose of 0.75 mg/kg, was either applied ‘early’, i.e. together with the proconvulsant 0 Mg^2+^ solution (Fig. 2C, Supp. Fig. 2H), or ‘late’ once epileptiform activity had already entered the SE-like late recurrent discharge (LRD) phase (Fig. 2L, Supp. Fig. 2P). Early application of diazepam significantly delayed the onset of SLEs as compared to control slices, which were not exposed to diazepam prior to onset of SLEs (Fig. 2E, Supp. Fig. 2K). This effect was particularly pronounced in acute slices where early application of diazepam often prevented SLE generation compared to control slices (Supp. Fig. 2I). Early application of diazepam also caused a significant decline in the total number of SLEs per slice (Fig. 2F, Supp. Fig. 2L) as compared to control. Moreover, early application of diazepam significantly delayed the onset of LRD in the organotypic brain slices (Fig. 2G) as well as the time between the 1^st^ SLE and LRD onset (Fig. 2H). The antiseizure effects of diazepam appeared more pronounced in the acute slices where the early diazepam significantly decreased the propensity for SLEs and LRD to (Supp. Fig. 2I and J). This demonstrated that diazepam had a significant antiseizure effect on the initial pathological discharges in 0 Mg^2+^.

In contrast, we found that in both organotypic and acute brain slices, diazepam lost its antiseizure effect once epileptiform activity had become persistent (SE-like activity) and instead enhanced discharges (Fig. 2I-P, Supp. Fig. 2M-T). To quantify this we measured the power spectral density (PSD) in a 5 min window 1-2 mins after the onset of LRD (‘Pre-window’, Fig. 2J, M and Supp. Fig. 2N, Q) as well as the PSD in a 5 min window 10-15 mins following the application of diazepam (‘Post-window’, Fig. 2K, N and Supp. Fig. 2O, R). In control slices diazepam was not present in the ‘Post-window’, but this window was taken at an equivalent time period after the ‘Pre-window’ and onset of LRD. This allowed us to calculate the change in power for each slice (‘Post-win. power’ - ‘Pre-win. power’ / by ‘Pre-win. power’) which was then normalised by the mean change in power from control slices to generate a metric we term the normalised change in power. We found that in the frequency range 1-40 Hz, diazepam application during LRD significantly increased the normalised change in power as compared to controls in both organotypic (2O) and acute (Supp. Fig. 2S) slices. Diazepam did not significantly affect the normalised change in power in the 60-90 Hz frequency range (Fig. 2P and Supp. Fig. 2T). Together, this demonstrates that ‘late’ diazepam application during LRD exacerbates epileptiform activity in two different brain slice preparations.

Phenobarbital, a barbituarate, is also a positive modulator of GABA_A_Rs and is sometimes used as second-line management for paediatric CSE refractory to benzodiazepines (Burman *et al*., 2019). In addition at higher concentration, it has been shown to antagonise AMPA/ Kainate receptors (Nardou *et al*., 2011). We therefore sought to determine the effect of both a low concentration of phenobarbital (100μM, equivalent to a total clinical dose of 16 mg/kg) and a high concentration of phenobarbital (300μM, equivalent to a total clinical dose of 49 mg/kg) on late stage SE-like activity in organotypic brain slices (Supp. Fig. 3). Strikingly, we found that like diazepam, 100μM phenobarbital significantly increased the normalised change in power (1-40 Hz) compared to controls (Supp. Fig. 3, A, B,C,G). In contrast, 300μM phenobarbital attenuated SE-like activity, significantly reducing the normalised change in power (1-40 Hz) compared to controls (Supp. Fig. 3, D, E, F, G). No significant changes were observed in the 60-90 Hz frequency range.

### Persistent epileptiform activity is associated with a reduction in GABAergic synaptic conductances

Having observed that diazepam and low dose phenobarbital lost their antiseizure efficacy during LRD, we next sought to determine whether synaptic GABA_A_R internalisation (Goodkin et al., 2005) and hence a reduction in GABA synaptic conductance (g_GABA_) could underlie this effect.

We optogenetically activated GABAergic neurons, and recorded postsynaptic GABA-R mediated currents, in voltage-clamp mode, in CA1 pyramidal neurons in mouse organotypic slice cultures. (Fig. 3A). By recording GABA currents at different holding voltages g_GABA_ could be measured before and after 30 mins exposure to 0 Mg^2+^ and confirmation of LRD in the same neuron. GABA_B_ receptor blockade to isolate GABA_A_Rs was not utilised as this could interfere with the seizure-like activity evoked by 0 Mg^2+^ (Swartzwelder *et al*., 1987; Codadu *et al*., 2018). Following termination of SE-like activity using reintroduction of Mg^2+^, g_GABA_ was remeasured. The mean g_GABA_ elicited under baseline conditions decreased after SE-like activity had been arrested (Fig. 3B-D), with no significant change in access resistance (baseline: mean 15.68 ± *SEM* 1.03MΩ vs 30 min after 0 Mg^2+^: mean 18.60 ± *SEM* 1.18MΩ, *p* =0.11, *paired t-test, data not shown*). These findings demonstrate that SE-like activity is associated with reductions in GABA synaptic conductances which is most likely caused by GABA_A_R internalisation (Goodkin et al., 2005). Nonetheless, despite prolonged periods of persistent seizure-like activity *in vitro*, optogenetic activation of GABAergic interneurons could still reliably evoke GABA synaptic currents.

**Figure 3:**
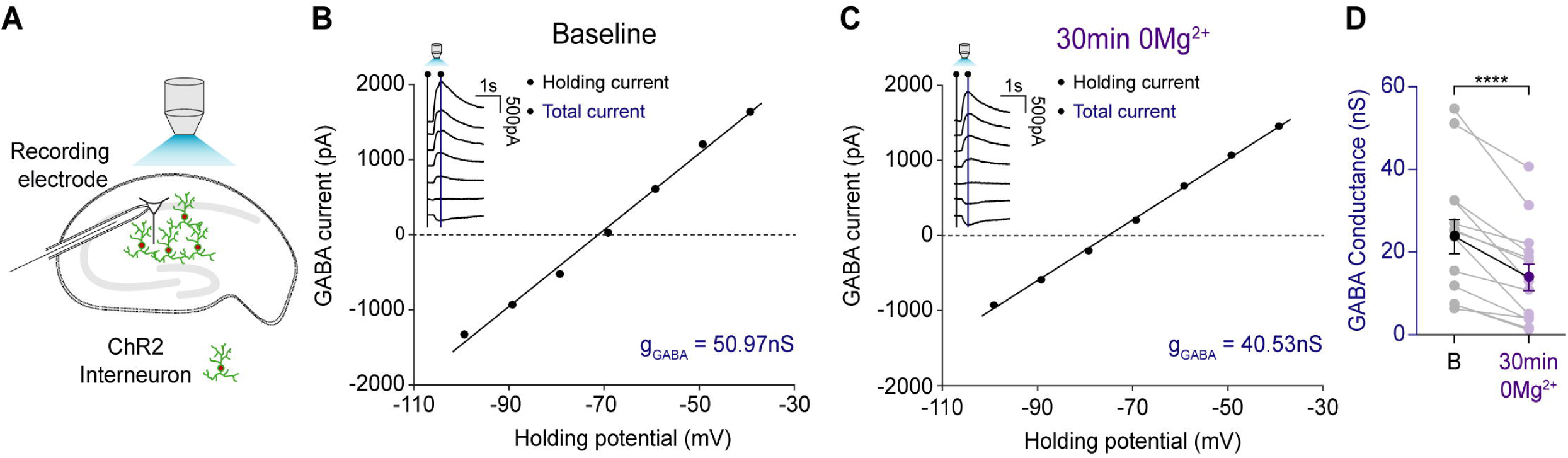
Persistent epileptiform activity is associated with a reduction in GABAergic synaptic conductance. **A**, Schematic of experimental setup showing whole-cell recordings being performed from CA1 pyramidal neurons in mouse organotypic brain slices where GAD2+ interneurons were transfected with ChR2-YFP and activated using a high-powered LED coupled to the objective. **B**, Example GABA I-V plot from a whole-cell patch-clamp recording (low Cl^-^ internal, 10 mM) of a CA1 pyramidal cell in a mouse hippocampal organotypic slice. Inset, raw current traces recorded at different holding potentials. GABA were evoked by optogenetic activation of ChR2 expressing GAD2+ interneurons with 100 ms blue light pulses. GABA conductance (g_GABA_) was calculated from the slope of the GABA current I-V curve. The GABA current was calculated by subtracting the holding current (black line on inset) from the total current (blue line on inset) for each holding potential. **C**, Example GABA I-V plot from the same cell as in ‘B’ following cessation of persistent epileptiform activity generated by 30 min of 0 Mg^2+^ application. **D**, Population (*n* = 14) data showing a significant decrease in g_GABA_ from baseline to after 30 min 0 Mg^2+^ (baseline: mean 23.6 ± *SEM* 4.1 nS vs 30min of 0 Mg^2+^: mean 13.1 ± *SEM* 3.2 nS, *p* = 0.0001, *paired t-test,*). \*\*\*\**p* ≤ 0.0001; error bars indicate mean ± SEM.

### Status epilepticus-like activity is accompanied by compromised neuronal Cl^-^ extrusion

We surmised that additional mechanisms to GABA_A_R internalisation must also play a role in the loss of diazepam’s antiseizure efficacy during LRD. This is because GABA_A_R internalisation cannot explain how diazepam and low dose phenobarbital could exacerbate epileptiform discharges during SE-like activity. Intracellular Cl^-^ accumulation and a depolarizing shift in E_GABA_, could potentially explain this phenomenon. To explore whether SE-like activity might compromise Cl^-^ extrusion mechanisms in neurons we performed gramicidin perforated patch-clamp recordings from hippocampal pyramidal cells. This technique avoids disrupting the [Cl^-^]_i_ of the neuron. Somatic application of GABA agonists (GABA, or muscimol) was used to record E_GABA_ under quiescent network conditions in order to assess steady state changes in Cl^-^ extrusion (Fig. 4A). E_GABA_ was measured using voltage step protocols, before Mg^2+^ removal (baseline) and after different periods of induced SE-like activity had been arrested by reintroducing Mg^2+^. We found that in the hippocampal organotypic brain slices, resting E_GABA_ became more depolarised after a period of 30 min of Mg^2+^ withdrawal (Fig. 4B-D). We further investigated this effect in rat hippocampal organotypic slices, extending the period of 0 Mg^2+^ withdrawal and putative SE-like activity to 3 hours. This also resulted in E_GABA_ becoming more depolarised (Fig. 4E).

**Figure 4:**
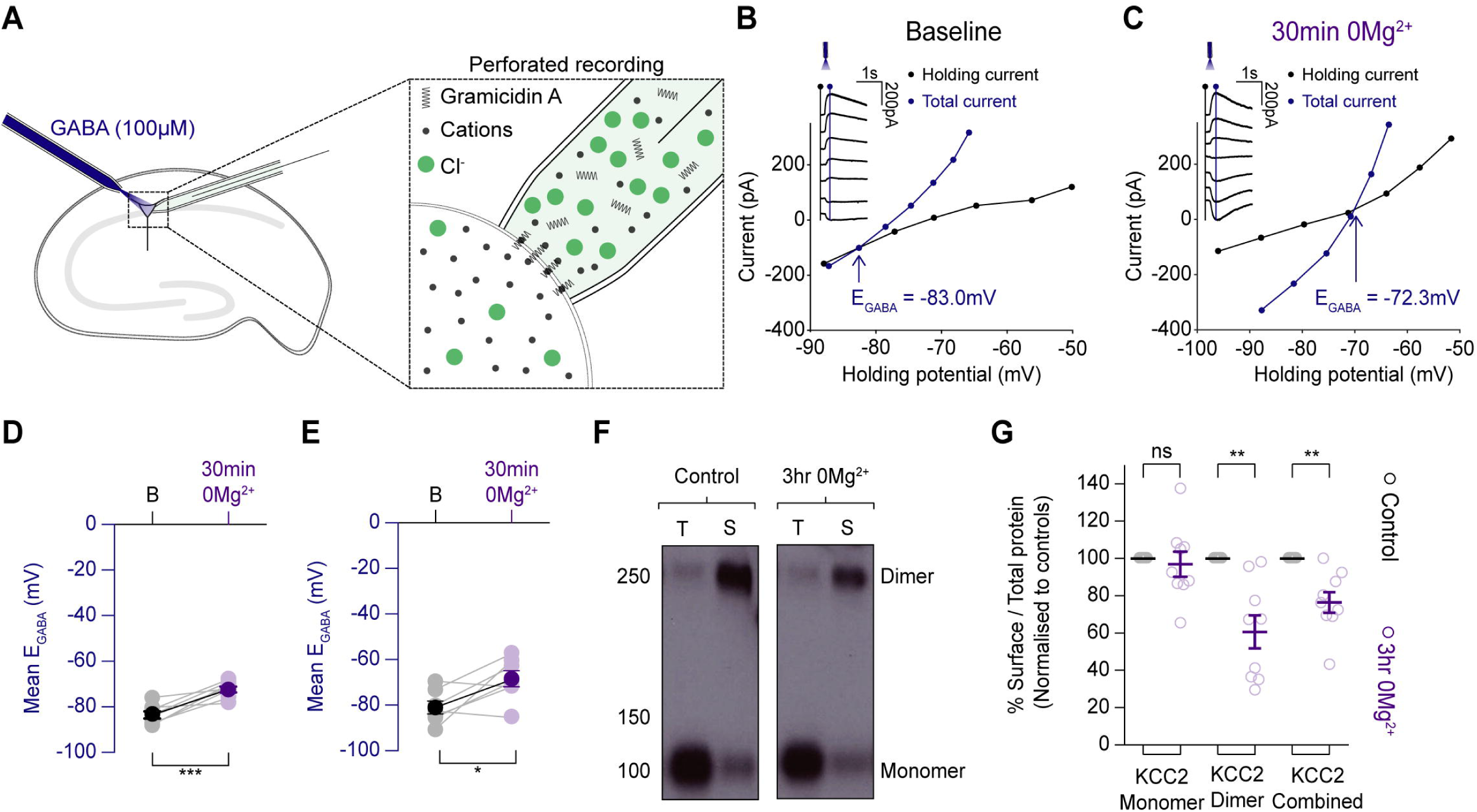
SE-like activity is associated with compromised neuronal Cl^-^ extrusion. **A**, Schematic demonstrating the gramicidin perforated patch-clamp recording configuration from organotypic hippocampal pyramidal cells and accompanying somatic GABA application. **B**, Example recording from a mouse CA1 pyramidal neuron. Resting E_GABA_ was measured by delivering GABA puffs at different holding potentials (inset). I–V curves were then plotted featuring the holding current (reflecting membrane current, black) and total current (reflecting membrane current plus the GABA-evoked current, blue). E_GABA_ was calculated as the potential at which the total current (blue line) was equal to the holding current (black line). **C,** Following 30 mins of Mg^2+^ withdrawal, epileptiform activity was arrested by reintroducing Mg^2+^ into the aCSF and resting E_GABA_ was once again measured as in ‘**B**’. **D**, Population data (*n* = 8) showing a significant increase in E_GABA_ between baseline and after a period of persistent seizure-like activity induced by 30 minutes of 0 Mg^2+^ application (resting: mean −83.4 ± *SEM* 1.5 mV vs after 30min 0Mg^2+^: mean −72.3 ± *SEM* 1.2 mV, *p* = 0.0008, *paired t-test*). **E,** Population data (*n* = 7) from a similar experiment as in ‘**F**’ from rat organotypic brain slices cultures and 3 hours of Mg^2+^ withdrawal demonstrating a further increase in E_GABA_ following this extended period of persistent epileptiform activity (resting: mean −80.8 ± *SEM* 2.8 mV vs after 3hr 0 Mg^2+^: −68.6 ± *SEM* 3.5 mV, *p* = 0.03, *paired t-test*). **F**, Western blots of control hippocampal slices, and those that had been treated with 3 hours of 0 Mg^2+^. Cell homogenates (‘T’-total) and NeutrAvidin captured cell surface proteins (‘S’-surface) were probed on western blots with the anti-C-terminus KCC2 antibody. 0 Mg^2+^ blots showed weaker bands for surface-bound KCC2 dimers, but little change in monomers compared to controls. **G**, Surface proteins from hippocampal slice lysates were quantified by the ratio of surface to total optical density. Values were then normalised to a percentage of untreated control values. Three hours of 0 Mg^2+^ treatment significantly reduced the surface: total ratio of overall KCC2 as compared to controls (surface KCC2 reduction, *n* = 9: 76.4 ± *SEM* 0.05% of control, *p* = 0.003, *unpaired t-test*) This change reflected a specific reduction in the surface levels of KCC2 dimers (surface KCC2 dimer reduction: 60.6 ± *SEM* 8.8% of control, *p* = 0.002, *unpaired t-test*). In contrast, monomeric KCC2 showed little difference across the two conditions with the mean in the 0 Mg^2+^ condition at (reduction of KCC2 monomer: 97.0 ± *SEM* 6.7% of control, *p* = 0.66, *unpaired t-test*) \**p* ≤ 0.05; \*\**p* ≤ 0.01; ‘ns’; not significant (*p* ≥ 0.05).

We next used surface biotinylation and western blotting for the major cation-chloride cotransporter (KCC2) to determine whether the depolarizing shift in E_GABA_ was accompanied by a shift in the cell surface expression of KCC2. Indeed, on average, the total amount of surface KCC2 in organotypic hippocampal slices treated with 0 Mg^2+^ was reduced to mean 76.4 ± *SEM* 0.05% of that found in control slices (Fig. 4F,G). When KCC2 was further subdivided into monomeric and dimeric forms it was found that this drop-in surface protein was almost exclusively due to a reduction in the KCC2 dimer. Zero Mg^2+^ treatment reduced the levels of surface bound KCC2 dimer significantly compared to controls. These findings confirm previous reports (Rivera *et al*., 2004) that prolonged seizure activity results in a reduction in expression of KCC2 and a depolarizing shift in steady-state E_GABA_.

### Persistent epileptiform activity drives pronounced depolarizing shifts in E_GABA_ and intracellular Cl^-^ accumulation

The shifts in steady-state E_GABA_ we observed above are unlikely to explain the enhancing effect of diazepam on burst discharges we observed during SE-like activity. This is because resting E GABA^’^s of ∼ −60 mV are typically below the action potential (AP) threshold from of CA1 neurons in our preparation (AP threshold: mean −38.72 ± *SEM* 5.45mV, *n* = 17, *data not shown*) and thus would still render GABA_A_R mediated transmission inhibitory. We therefore surmised that compromised Cl^-^ extrusion combined with the activity-dependent Cl^-^ loading during SE-like activity could result in more severe shifts in E_GABA_.

To explore this possibility, we modified our gramicidin perforated patch-clamp recording protocols in order to track dynamic changes in E_GABA_ during the evolution of epileptiform activity in the 0 Mg^2+^ model (Fig. 5A). The recording configuration was rapidly switched between current-clamp mode in order to measure membrane potential and short periods in voltage-clamp mode, during which a voltage ramp protocol and somatic GABA puff was applied, to provide a rapid estimate of E_GABA_ (Fig. 5B). Compared to baseline, SLEs and SE-like activity (LRD phase) were associated with pronounced increases in E_GABA_ (Fig. 5C,D). Once persistent epileptiform activity was terminated by tetrodotoxin (TTX), E_GABA_ decreased but did not return to baseline levels. The E_GABA_ remained moderately but persistently depolarised, further confirming the long-term shifts in Cl^-^ extrusion mechanisms described above. These experiments demonstrated E_GABA_ to be highly dynamic. SE-like activity was associated with profound elevations in E_GABA_ to values comparable to the AP threshold for these neurons, which would render GABA_A_R mediated transmission excitatory.

**Figure 5:**
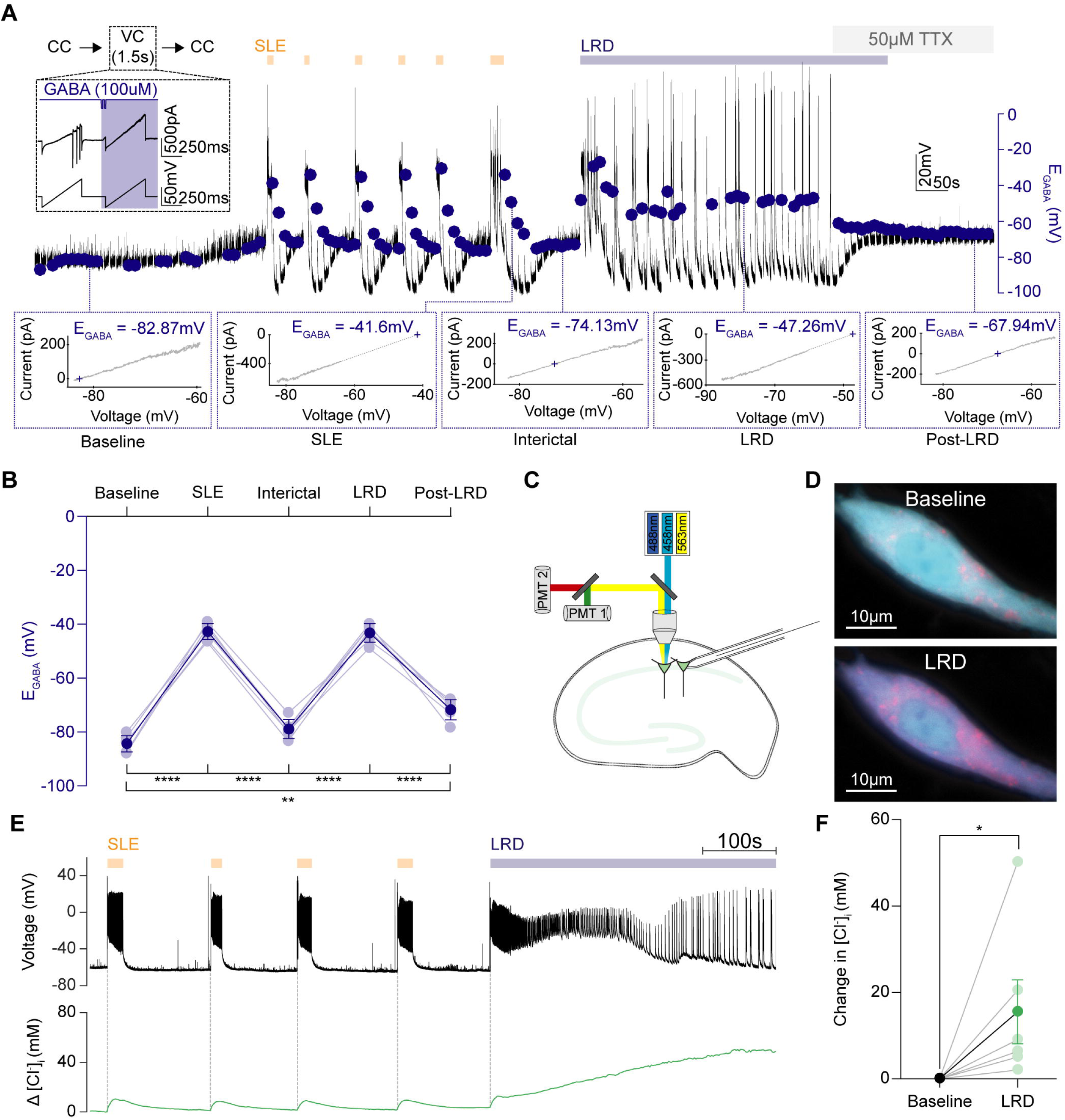
Persistent seizure-like activity drives pronounced depolarizing shifts in E_GABA_ and intracellular Cl^-^ accumulation. **A**, To measure E_GABA_ during epileptiform activity, gramicidin perforated patch-clamp recordings were performed and the recording mode was rapidly switched from current-clamp (CC) to brief periods in voltage clamp (1.5 s duration) every 10 s (inset). While in voltage-clamp, two consecutive voltage ramps were applied: (1) the first without GABA application; and the second paired with GABA application (blue) applied to the soma. A representative recording from a CA1 pyramidal neuron where E_GABA_ measurements (blue dots) were made throughout the progression of epileptiform activity in the 0 Mg^2+^ model (orange arrows shows SLEs, blue bar depicts LRD). Dotted lines highlight periods during evolution of epileptiform activity: baseline (*t_1_*), immediately following SLEs (*t_2_*), between SLEs / interictal (*t_3_*), LRD (*t_4_*), following termination of activity / post-LRD (*t_5_*). To rapidly abort epileptiform activity during LRD, TTX (50 μM) was applied. Bottom, I-V plots were used to calculate E_GABA_ defined as the voltage at which the GABA current equals 0 (*t_1-5_*). **B**, Population data showing significant changes in E_GABA_ between the different periods. E_GABA_ shifted from mean baseline levels of mean −83.9 ± *SEM* 1.1 mV to mean - 42.7 ± *SEM* 1.0 mV during SLEs (*n* = 7, *p* < 0.0001, *paired t-test*) before partially recovering to a mean level of mean −78.9 ± *SEM* 1.1 mV between events (*p* < 0.0001, *paired t-test*). SE-like activity (LRD phase) profoundly elevated E_GABA_ again to mean −43.2 ± *SEM* 1.9 mV (*p* < 0.0001 as compared to baseline, *paired t-test*). E_GABA_ levels were equally high during SLEs and the LRD phase (SLE: mean −42.7 ± *SEM* 1.04 mV vs LRD: mean −43.2 ± *SEM* 1.9 mV, *p* = 0.73, *paired t-test*). Following termination of the LRD phase with TTX, the EGABA decreased but remained more depolarised compared to the baseline E_GABA_ (baseline: mean - 83.9 ± *SEM* 1.13mV vs recovery: mean −71.0 ± *SEM* 1.5mV, *p* = 0.001, *paired t-test*). **C**, A schematic showing the experimental setup for Cl^-^ imaging in which hippocampal pyramidal neurons expressing ClopHensorN were imaged, while a simultaneous patch-clamp recording was performed from a neighbouring neuron. To determine [Cl^−^]_i_, confocal images were collected following excitation at 458, 488, and 563nm, respectively. **D**, Confocal images of the neuron in ‘E’ with the 458 nm and 563 nm fluorescence emission channels superimposed during baseline (top) and LRD (bottom). The fluorescence ratio from these channels (F458/F563) is sensitive to [Cl^−^]_i_, hence the shift to pink during LRD indicates an increase in [Cl^−^]_i_. **E**, Simultaneous measurement of activity-dependent changes in [Cl^−^]_i_ in a CA1 hippocampal pyramidal neuron expressing ClopHensorN (green trace, bottom). A current-clamp recording from a neighbouring pyramidal neuron (black trace, top; cell somata <200µm apart) provided a readout of epileptiform activity, including SLEs (orange arrows) and LRD (blue bar). **F**, Population data (*n* = 6) showing significant increases in [Cl^-^]_i_ associated with LRD compared to baseline activity (increase in [Cl^-^]_i_: mean 15.53 ± *SEM* 7.39 mM, *p* = 0.03, *Wilcoxon test*). \**p* ≤0.05; \*\**p* ≤0.01; \*\*\*\**p* ≤0.0001; error bars indicate mean ± SEM.

To directly measure changes in Cl^-^ concentration, which could underlie the observed activity driven shifts in E_GABA_, we employed the genetically encoded reporter of Cl^-^, ClophensorN. This reporter enables pH corrected estimates of intracellular Cl^-^ concentration (Arosio *et al*., 2010; Raimondo *et al*., 2013; Sato *et al*., 2017). Biolistic transfection of mouse organotypic hippocampal brain slices resulted in sparse ClopHensorN expression within pyramidal neurons. Confocal imaging of ClopHensorN expressing cells was performed concurrently with whole cell patch-clamp recordings of neighbouring cells to provide a simultaneous readout of seizure-like activity (Fig. 5C-E). SLEs and the LRD phase were associated with increases in [Cl^-^]_i_ (Fig. 5E, F). Importantly, the LRD phase was associated with a significant increase in [Cl^-^]_i_. These results suggest that *in vitro* SE-like activity is accompanied by profound short-term, activity-driven increases in E_GABA_ and [Cl^-^]_i._

To determine whether the canonically inward Cl-cotransporter (NKCC1), might contribute to Cl-accumulation and the effects of diazepam on late stage SE-like activity, we repeated the interface experiments in Fig. 2. I-Q, but combined diazepam application with 10μM bumetanide to block NKCC1 activity (Supp. Fig. 4). We found that the addition of bumetanide did not significantly alter the normalised change in power as compared to control slices or to diazepam application alone (Supp. Fig. 4A-E). This result suggests that blocking NKCC1 during LRD does not rescue the antiseizure effects of diazepam during this phase of epileptiform activity.

### GABA-releasing interneurons are active and highly correlated with pyramidal cell activity during the late recurrent discharge phase

Having observed significant increases in E_GABA_ and intracellular Cl^-^ in pyramidal neurons during the LRD phase of the 0 Mg^2+^ model of SE, we next aimed to determine how the activity of GABA-releasing interneurons might relate to that of pyramidal cells during the recurrent discharges observed in this period. To do so we used organotypic slices prepared from mice where the cre-lox system was used to selectively express the red fluorescent reporter tdTomato under the glutamic acid decarboxylase type 2 (GAD2) promoter. Using dual whole-cell patch-clamp, we made targeted recordings from CA1 hippocampal GABAergic interneurons and pyramidal cells during various phases of seizure-like activity in the 0 Mg^2+^ *in vitro* model of SE (Fig. 6A). We noticed that GABAergic interneurons were highly active during the LRD phase (Fig6. C, D). To determine whether the activity of GABAergic interneurons was synchronised with that of pyramidal cells we used linear correlation as a measure of synchrony (Jiruska *et al*., 2013). We compared the synchrony during baseline, single SLEs, during the LRD phase and following cessation of epileptiform activity (post-LRD). The correlation and hence synchrony between interneurons and pyramidal cells was increased during SLEs and LRD as compared to pre and post epileptiform activity (Fig6. D, E). Synchrony between these two cell types was significantly higher during LRD as compared to SLEs, which demonstrates that persistent epileptiform activity is composed of highly synchronous activity between GABAergic interneurons and glutamatergic pyramidal cells.

**Figure 6:**
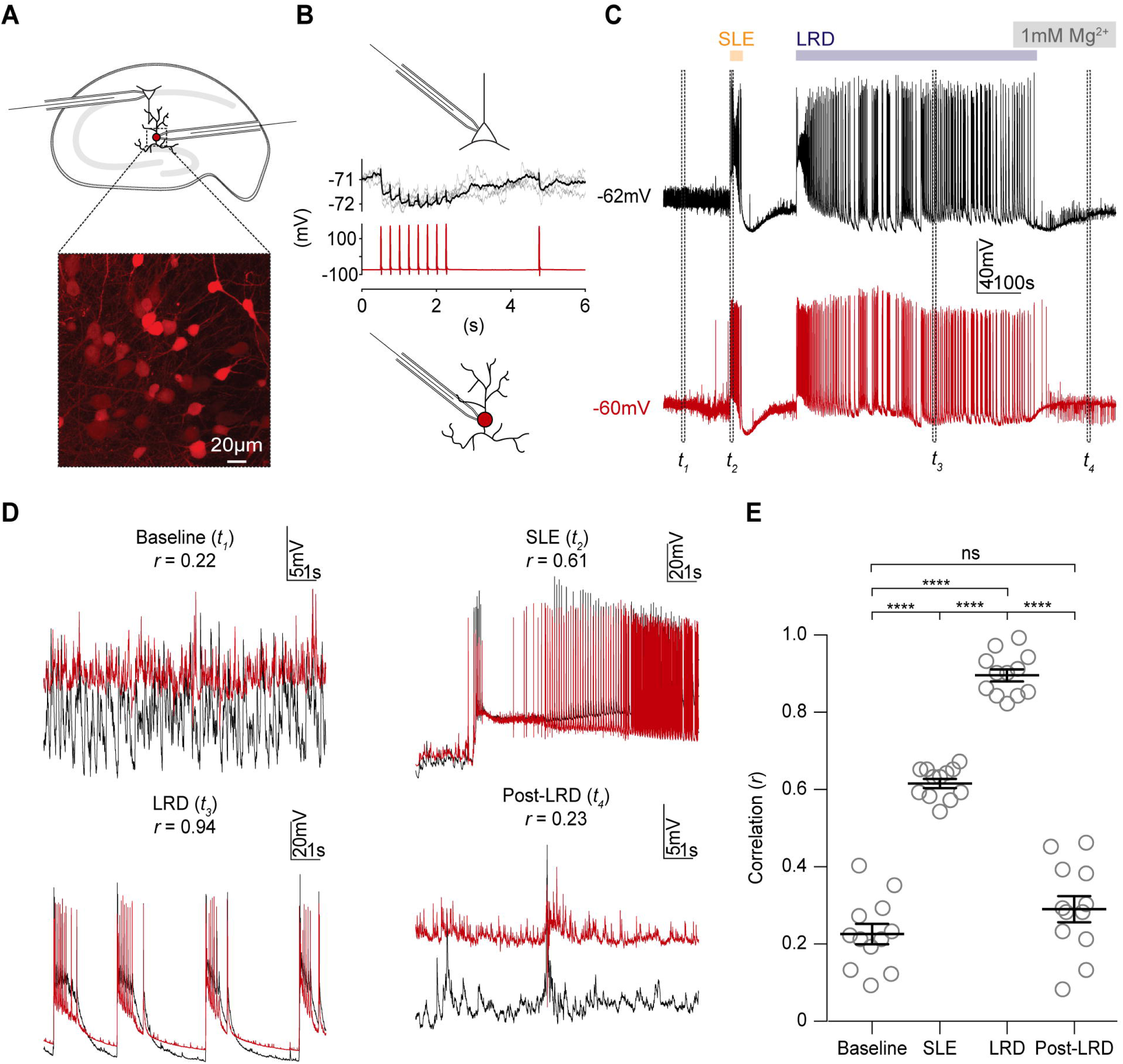
GABA-releasing interneurons are active and highly correlated with pyramidal cell activity during the late recurrent discharge phase. **A**, Diagram of the experimental setup (top), which involved simultaneous whole-cell patch-clamp recordings from CA1 pyramidal neurons and GABAergic interneurons in organotypic hippocampal brain slices (*n* = 12). Inset, confocal images show interneurons in the stratum radiatum expressing tdTomato using the cre-lox system (tdTomato reporter line crossed with GAD2-cre line). **B**, GABAergic connection between pyramidal cell (black) and GAD2+ interneuron (red) confirmed by observing negative shifts in the pyramidal cell membrane potential when action potentials were generated in the interneuron. Only 3 of the 12 paired recordings showed functional inhibitory synaptic connections. **C**, Dual current-clamp recordings from the same cells in ‘**B**’ during progression of 0 Mg^2+^ seizure-like activity. Four periods are denoted by dashed rectangles: baseline (*t_1_*), at the start of a single SLE (*t_2_*), during the LRD phase (*t_3_*) and after epileptiform activity had been aborted with the re-introduction of 1 mM Mg^2+^, post-LRD (*t_4_*). **D**, Insets show the four periods in ‘**C**’, with recordings from the two neurons superimposed to reveal the extent of synchronous activity. The Pearson’s coefficient (*r*) was calculated from a direct linear correlation of the two raw traces during each epoch (8 s duration). **E**, Population data showing significant increases in correlation during single SLEs (*t_2_*) and the LRD phase (*t_4_*) (baseline mean *r* = 0.2 ± *SEM* 0.03; SLEs mean *r* = 0.6 ± *SEM* 0.01; LRD mean *r* = 0.90 ± *SEM* 0.02, post mean *r* = 0.29 ± *SEM* 0.03, *p* < 0.0001, *paired t-tests,*). Furthermore, the synchrony between pyramidal cells and interneurons was greater during LRD compared to during single SLEs (mean *r* = 0.9 vs mean *r* = 0.6, *p* < 0.0001, *paired t-test*). ‘GAD2’, glutamic acid decarboxylase 2; ‘ns’, non-significant; ‘SR’, stratum radiatum; ‘Td’, tandem dimeric tomato. \*\*\*\**p* ≤ 0.0001; ‘ns’, not significant (*p* ≥ 0.05); error bars indicate mean ± SEM.

### GABAergic signalling is strongly depolarising during the late recurrent discharge phase

Given our observations of a depolarizing E_GABA_ and highly synchronized GABAergic interneuronal and pyramidal cell activity during the SE-like activity of the LRD phase, we next explored whether GABAergic signalling might in fact be excitatory during LRD. To investigate this, we used an optogenetic approach to isolate and selectively activate ChR2-expressing GABAergic (GAD2+) interneurons during different phases of seizure-like activity. Using this experimental setup, GABAergic interneurons were activated every 7 s with 100 ms of blue light whilst performing whole-cell current-clamp recordings from CA1 pyramidal neurons during the progression of seizure-like activity in the 0 Mg^2+^ model.

We observed that whilst light activation resulted in hyperpolarizing synaptic potentials under baseline conditions, (Fig. 7A-C), during single SLEs, light delivery reliably triggered membrane depolarisation and the generation of action potentials (Fig. 7B, C). In-between SLEs, the hyperpolarising responses to optogenetic activation were restored. However, during the SE-like LRD phase, light activation again consistently resulted in strong depolarisation and action potentials in the recorded neurons (Fig. 7C).

**Figure 7:**
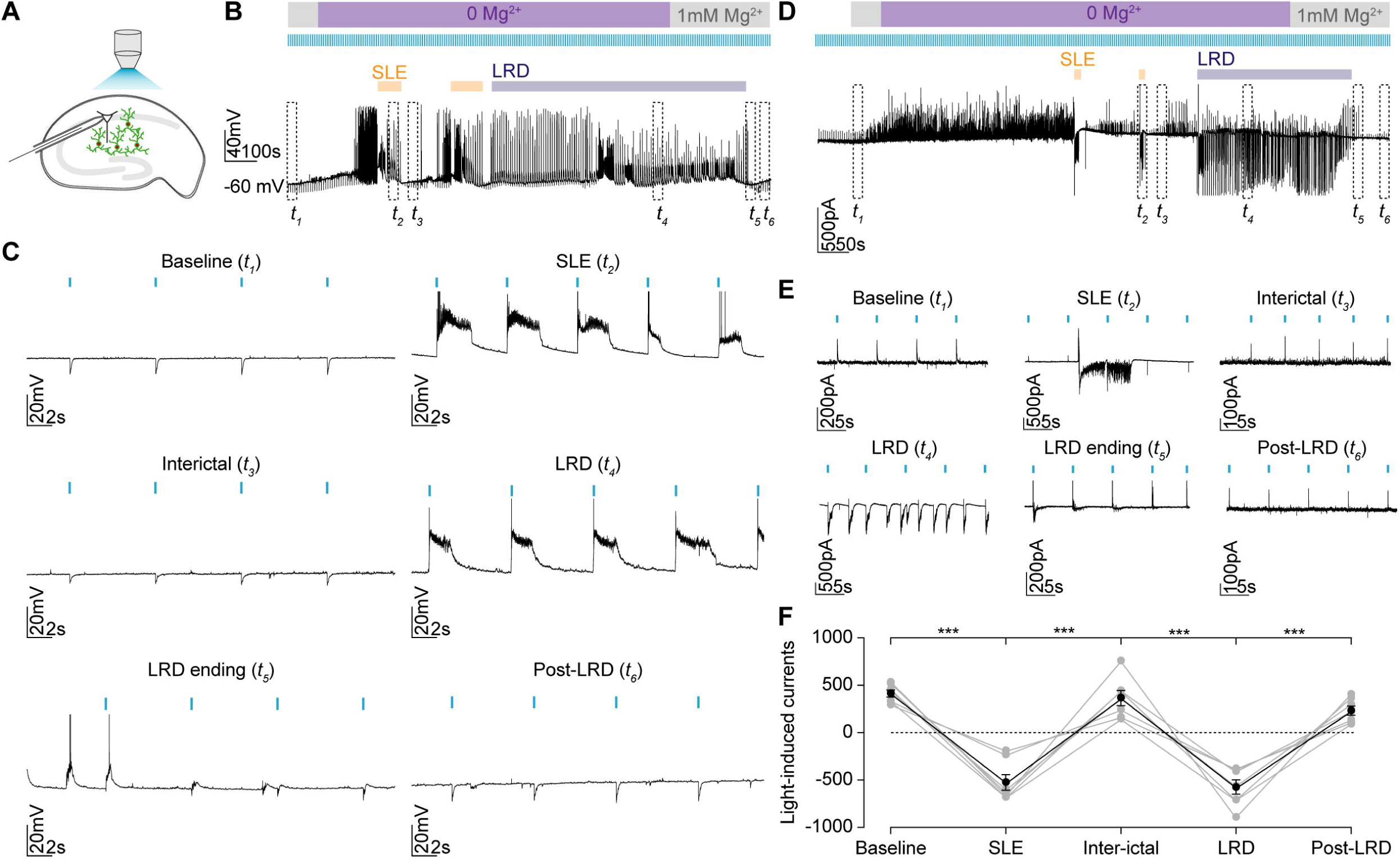
GABAergic signalling is strongly depolarising during the late recurrent discharge phase. **A**, Experimental setup showing whole-cell recordings being performed from CA1 pyramidal neurons in mouse organotypic brain slices where GAD2+ interneurons were transfected with ChR2-YFP and activated using a high-powered LED coupled to the objective. **B**, Current-clamp recording with optogenetic activation of GAD2+ interneurons every 7s using 100 ms light pulses during the evolution of epileptiform activity in Mg^2+^-free solution. Individual SLEs (orange bars) and LRD are indicated (blue bar). The persistent activity of the LRD phase was terminated using aCSF containing 1mM Mg^2+^. The dashed rectangles *t*_1-6_ represent 30 s windows from different periods: baseline (*t*_1_), latter portion of a SLE (*t*_2_), interictal period (*t*_3_), LRD (*t*_4_), LRD ending (*t*_5_) and post-LRD (*t*_6_). **C**, Expanded view of the windows *t*_1-6_ in ‘**B**’ show optogenetic activation of GAD2+ interneurons (blue bars) shifting from causing membrane hyperpolarisation during baseline and interictal periods to membrane depolarisation and action potentials during SLEs and LRD. These effects were transient with the cessation of LRD resulting in a return of IPSPs. **D**, Voltage-clamp recording from CA1 pyramidal cell clamped at −40 mV to record light-induced currents using the protocol as in ‘**B**’. Dashed rectangles representing the same periods as in ‘**B**’. **E**, Expanded views of *t*_1-6_ showing changes in the maximum light-induced currents at each phase of activity. **F**, Population data (*n* = 7) showing significant changes in current size and direction across the different phases. Light currents were recorded as the maximum light induced current within 100 ms of the light pulse. At rest these were outward (positive), with a mean value of mean 448.8 ± *SEM* 70.3 pA but rapidly flipped to being inward (negative) during SLEs (mean −752 ± *SEM* 88.7 pA, *p* = 0.0002, *paired t-test*). After the SLE had recovered, the currents returned to positive values (mean 536.5 ± *SEM* 134.9 pA, *p* = 0.0007). However, during the LRD phase the currents again became significantly negative compared to the preceding interictal period (mean −894.0 ± *SEM* 164.8 pA, *p* = 0.0018, *paired t-test*). When Mg was reintroduced, the current again shifted to become positive (mean 242.0 ± *SEM* 21.2 pA, *p* = 0.0005, *paired t-test*). There was no significant difference in amplitude between light-induced currents during SLE and LRD (SLE: mean −752.00 ± *SEM* 88.67 pA vs LRD: mean −894.00 ± *SEM* 164.80 pA, *p* = 0.18, *paired t-test*). \*\*\**p* ≤ 0.001; error bars indicate mean ± SEM.

We then repeated this with cells held at −40 mV in the voltage-clamp configuration in order to record light-induced synaptic currents (Fig. 7D-F). After SLEs had self-terminated, light-induced currents returned to positive values. During the LRD phase, the currents recorded in pyramidal neurons following optogenetic activation of GABAergic interneurons again became significantly negative. This could be reversed when persistent epileptiform activity was terminated using reintroduction of Mg^2+^-containing aCSF. Taken together, this data demonstrated that GABAergic interneurons have profound excitatory effects on their synaptic targets during persistent epileptiform activity in the 0 Mg^2+^ model.

### Optogenetic activation of GABAergic interneurons triggers epileptiform discharges and entrains the hippocampal network during persistent epileptiform activity in a *GABA_A_R* dependent manner

Our data suggests that the GABAergic inhibitory system becomes ineffective during SE-like activity due to changes in chloride homeostasis and this could explain the failure of diazepam to reduce seizure-like activity during this phase. To further test this hypothesis, we next sought to determine whether recruitment of GABAergic interneurons fails to inhibit epileptiform activity during LRD and furthermore, whether selective activation of interneurons results in more pro-seizure like effects than antiseizure activity as was witnessed following diazepam application in our preparation.

During a 5 min analysis period during the LRD phase, we analysed our traces in 7 s windows to determine what effect selective activation of GABAergic interneurons (100 ms blue light activation occurring 3.5 s into the 7 s window) had on epileptiform activity. This 7 s window was then divided into smaller 200 ms time bins whereby the probability of an epileptiform discharge being initiated could be calculated (Fig. 8A-D). Discharges were defined as inward currents greater than 10 standard deviations from baseline noise and lasting more than 500 ms. We observed that the probability of discharge initiation increased to mean 0.78 ± *SEM* 0.06 during the 200 ms immediately following light application. In a control experiment repetitive light activation was initiated only after a minimum of 5 min of LRD activity had occurred. The probability of discharge initiation within the 200 ms time bin 3.5 s into the 7 s window increased when the light was delivered at the start of this time bin (Fig. 8E-I).

**Figure 8:**
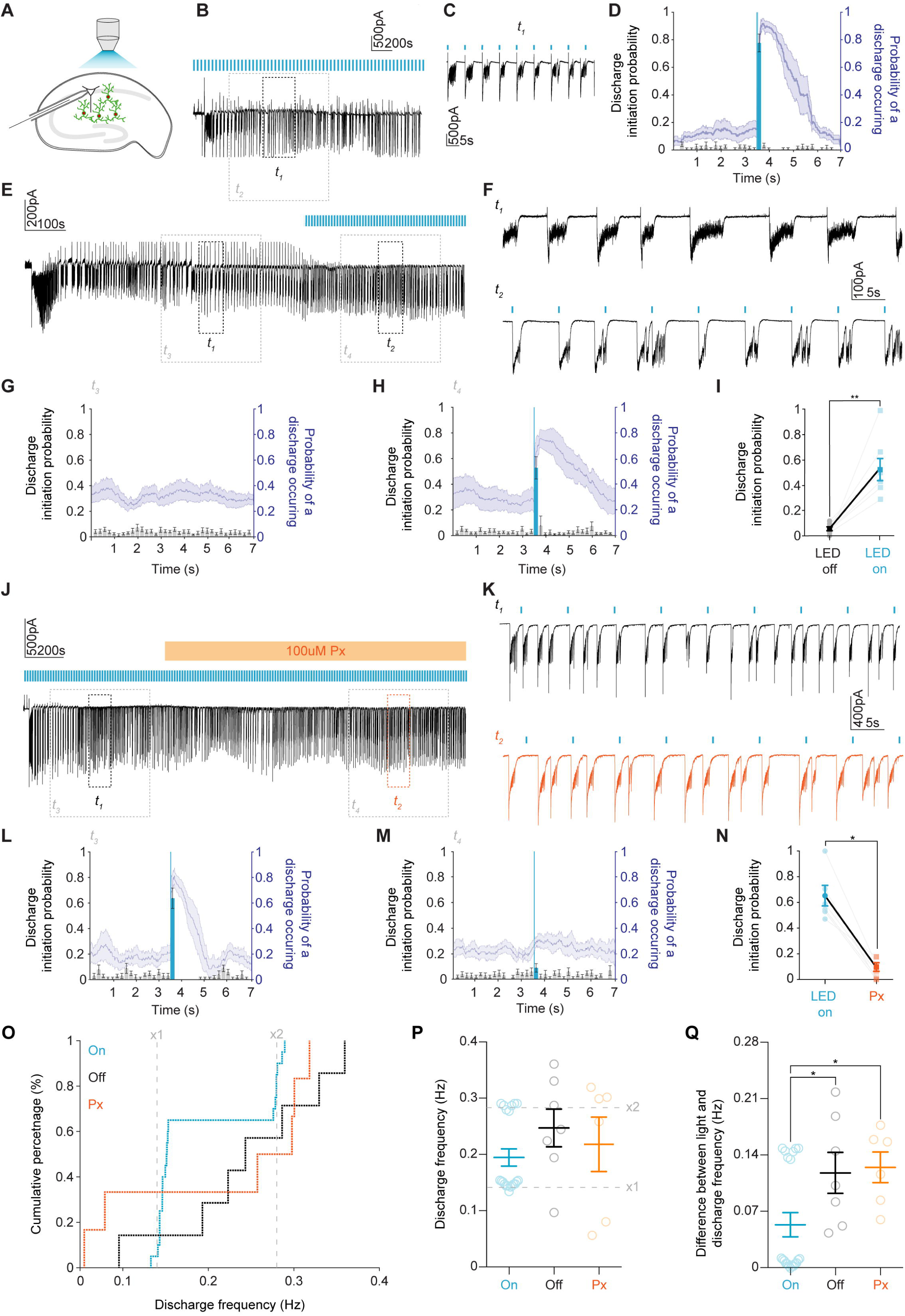
Optogenetic activation of GABAergic interneurons during LRD triggers burst discharges and entrains the hippocampal network in a GABA_A_R dependent manner. **A**, Experimental schematic showing whole-cell patch-clamp recordings from CA1 pyramidal neurons with widefield optogenetic activation of GAD2+ interneurons. **B**, Voltage-clamp recording during LRD from a cell clamped at −40 mV with 100 ms blue light pulses delivered once every 7 s. *t*_1_ is a 60 s window of activity which is expanded in **C** showing epileptiform discharges reliably initiated by optogenetic activation of GAD2+ interneurons. *t*_2_ (grey) demarcates section of trace used for analysis. **D**, Population data from 7 slices. Left axis (black, histograms with ± SEM) represents the probability of a discharge being initiated in any given 200 ms time bin within a 7 s window from a 5 min LRD analysis period. Light blue highlights the time bin where light was delivered 3.5 s into the 7 s window. The right axis (blue, line plot with ± SEM) represents the probability of a discharge being present. **E**, Trace showing a 10 min window from the onset of LRD. *t*_1_ and *t*_2_ are 60 second periods of activity before and after blue light was delivered as in ‘**B**’. **F**, Expanded *t*_1_ and *t*_2_ showing 100 ms blue light application (blue bars) reliably initiating discharges. *t*_3_ and *t*_4_ (grey) demarcates section of trace used for analysis. **G**, Population data (*n* = 7) as in ‘D’ from 5 min analysis periods during LRD prior to light activation. **H**, Same analysis as in ‘G’ for 5 min analysis periods from the same slices where 100 ms blue light was delivered 3.5 s into every 7 s window. **I**, Discharge probability significantly increased in the time window where blue light was applied (increased from mean 0.1 ± *SEM* 0.02 before the light pulses were delivered to mean 0.5 ± *SEM* 0.1 after light was delivered, *p* =0.002, *paired t-test*). **J**, 10-minute window from the onset of LRD with light-stimulus delivered every 7 seconds. After at least 5 minutes of LRD activity, picrotoxin (100 μM) was applied. *t*_1_ and *t*_2_ are 60 s periods of activity before and after picrotoxin (orange) application. **K**, Expanded views of *t*_1_ and *t*_2_ demonstrating that light delivery (blue bars) no longer elicited discharges. *t*_3_ and *t*_4_ (grey) demarcates section of trace used for analysis. **L**, Population data (*n* = 6) as in ‘**D**’ from the 5 min analysis window prior to picrotoxin application. **M**, Data from the same recordings as in ‘**L**’ for a 5 min analysis period in the presence of picrotoxin. **N**, Discharge probability significantly decreased in the time bin following blue light application in the presence of picrotoxin (decreased from median 0.6 *IQR* 0.5 – 0.8 to median 0.1 *IQR* 0.0 – 0.2 following GABA_A_R blockade with picrotoxin, *p* = 0.03, *Wilcoxon test,*). **O**, Cumulative percentage plot showing the distribution of discharge frequencies across three groups, control with photoactivation (‘On’, blue, *n* = 20), no photoactivation (‘Off’, *n* = 7) and photoactivation in the presence of picrotoxin (‘Px’, *n* = 6). Grey lines mark x1 and x2 the frequency of the light stimulus (0.14Hz and 0.28Hz). **P**, Population data of discharge frequencies. **Q**, The difference between frequency of photoactivation (x1 grey line in ‘**O’**) and the observed frequency of discharges within a 5 min analysis window. In the ‘On’ group, this difference was significantly smaller compared to the ‘Off’ and Px groups (‘On’: median 0.01 *IQR* 0.01 – 0.1Hz vs ‘Off’: median 0.1 *IQR* 0.1 – 0.2Hz, *p* = 0.03, *Mann-Whitney U test* and vs ‘Px’: median 0.1 IQR 0.1 – 0.2Hz, *p* = 0.01, *Mann-Whitney U test*). \**p* ≤ 0.05; \*\**p* ≤ 0.01; error bars indicate mean ± SEM.

Having established that optogenetic activation of GABAergic interneurons reliably initiated epileptiform discharges during SE-like activity, we next sought to determine whether excitatory GABA_A_R synaptic signalling was responsible for this effect. To do so we added the GABA_A_R antagonist picrotoxin (100 μM) to the perfusing aCSF after at least 5 minutes of LRD activity had been induced (Fig. 8J - N). The picrotoxin did not arrest the LRD activity demonstrating that GABA_A_R mediated synaptic transmission is not necessary for the generation of these discharges. However, we did note that the probability of discharge initiation following optogenetic activation of GABAergic interneurons decreased significantly (Fig. 8J - N). This confirmed that GABA_A_R mediated transmission is necessary for the pro-seizure properties of the GABAergic interneuronal network during LRD.

Finally, we measured changes in the frequency of epileptiform discharges during LRD when GABAergic interneurons were optogenetically activated at 0.14 Hz (‘On’), when the light was not applied (‘Off’) and when optogenetic activation occurred in the presence of 100 μM pictrotoxin (‘Px’). As demonstrated in Fig. 8O and P, in the ‘On’ condition, the frequency of discharges was entrained either to the frequency of light delivery (0.14 Hz) or twice the frequency (0.28Hz), indicating one ‘break through’ discharge every cycle. By comparison, if there was no light activation or picrotoxin was present, there was a wide distribution of discharge frequencies. In addition, there was a significantly smaller difference between the frequency at which the light was delivered and the frequency of discharges between the ‘On’ group as compared to the ‘Off’ (and picrotoxin groups (Fig. 8Q). Taken together this data demonstrates that GABAergic interneurons are ineffective at curtailing epileptiform discharges during SE-like activity and can enhance epileptiform activity.

## Discussion

In our study we have used both clinical and experimental data to explore the phenomenon of benzodiazepine-resistant SE. We found that the prevalence of benzodiazepine resistance in a South African cohort of paediatric patients in SE is similar to that reported previously in the UK (Chin *et al*., 2008). In addition, our clinical data confirm several prior observations, that provide some critical insights into the underlying pathology and clinical management. Firstly, benzodiazepine resistant SE is associated with increased morbidity in patients (Chin *et al*., 2008). Secondly, patients who seize for longer prior to initial treatment were more likely to be resistant to benzodiazepine treatment. While previous studies and reviews have alluded to longer SE duration being associated with increased resistance to benzodiazepines (Deeb *et al*., 2012; Naylor, 2014; Fernández *et al*., 2015; Gaínza-Lein *et al*., 2018; Gaínza-Lein *et al*., 2019), our clinical data are the first to quantify this phenomenon. Notably, we show that the duration of SE can successfully be used as a binary classifier to detect benzodiazepine resistance in SE. In our patient population, the optimum threshold for discriminating between benzodiazepine resistant versus sensitive patients was 61 mins of seizure activity.

Given the clinical importance of this phenomenon, and our observation that diverse brain insults causing prolonged seizures can all result in benzodiazepine resistance, we used the 0 Mg^2+^ *in vitro* model of SE to identify multiple changes in GABAergic signalling that could explain the progressive loss of diazepam efficacy in this condition, over this time frame. SE-like activity was accompanied by a modest reduction in GABA synaptic conductance and persistent changes in the efficacy of Cl^-^ extrusion. However, our major observation was profound short-term, activity-dependent Cl^-^ accumulation during SE-like activity. As a result, optogenetic activation of GABAergic interneurons was ineffective at reducing epileptiform activity; on the contrary, GABAergic interneuron firing actually enhanced epileptiform discharges during SE, via excitatory GABA_A_R mediated synaptic transmission. This activity-dependent effect is supplemental to two other changes we document, namely the reduced KCC2 expression, and the associated shift in baseline E_GABA_. Together, these results demonstrate why benzodiazepines may fail to enhance inhibition in continuously seizing brain circuits.

The withdrawal of Mg^2+^ *in vitro* has long been used as a model for studying putative changes in GABAergic signalling during SE (Dreier and Heinemann, 1991; Goodkin *et al*., 2005; Albus *et al*., 2008). Dreier *et al*. (1998) were the first to demonstrate that benzodiazepines lose their antiseizure efficacy following the onset of the SE-like LRD phase in the 0 Mg^2+^ model. We extend this work by demonstrating that during SE-like activity, not only do benzodiazepines lose their antiseizure action, but further, they can even exacerbate seizure-like activity. We suggest that the effects of diazepam during SE we observe are not explained by previously described deficits in GABA_A_R trafficking on principal cells alone (Goodkin *et al*., 2008), but also by a transient collapse in the transmembrane Cl^-^ gradient and EGABA.

Previous reports suggest that on-going seizure activity results in reduced surface expression, and function, of the canonical Cl^-^ extruder KCC2 (Rivera *et al*., 2004). There are multiple mechanisms by which this could occur including enhanced NMDAR activation, which is known to downregulate KCC2 function (Lee *et al*., 2011). This is unlikely to result purely from 0 Mg^2+^ treatment as prolonged seizure activity induced by 4-aminopyridine also reduces KCC2 function (Rivera *et al*., 2004). KCC2 ensures that Cl^-^ is maintained at levels lower than would be predicted by passive processes and also ameliorates activity-dependent Cl^-^ loading (Düsterwald *et al*., 2018). Loss of KCC2, therefore, constitutes a double blow to the system, since it will cause a rise in baseline intraneuronal [Cl^-^], and reduce the rate of clearance of Cl^-^, meaning that activity-dependent Cl^-^loading is exacerbated. We have shown both effects are important. Our findings confirm that the progression to SE-like activity in the 0 Mg^2+^ model is accompanied by a reduction in KCC2 surface expression and function as evidenced by a progressive depolarization of resting E_GABA_ prior to and following SE-like activity. In neonatal tissue on-going seizure activity is associated with enhanced activity of the Cl-importer NKCC1 (Dzhala *et al*., 2010). In our preparation, where Cl^-^ homeostasis mechanisms have matured, pharmacological blockade of NKCC1 did not appear to rescue the antiseizure effect of diazepam on SE-like activity. This suggests that NKCC1 is unlikely to contribute to the Cl^-^ loading we observe. This supports the importance of short-term, activity-dependent, increases in Cl^-^, likely via GABA_A_Rs themselves (Raimondo *et al*., 2015). These have previously been demonstrated during single SLEs both *in vitro* and *in vivo* (Isomura *et al*., 2003; Ellender *et al*., 2014; Sato *et al*., 2017), but our new data represent the first time this has been shown to be a factor in persistent SE-like activity. Our gramicidin perforated patch-clamp recordings and Cl^-^ imaging measurements using the genetically encoded ratiometric reporter ClopHensorN demonstrate that both single SLEs and SE-like activity result in substantial intracellular Cl^-^ accumulation and an excitatory shift in E_GABA_. Whether Cl-dysregulation itself drives the transition to SE-activity or is a product of SE-activity is still an open question. However, it is worth noting that reducing the function of KCC2 has been shown to accelerate the transition to SE both *in vitro* (Kelley *et al*., 2016) and *in vivo* (Silayeva *et al*., 2015).

Seizures are able to start and spread due to a loss of inhibitory synaptic mechanisms which are typically recruited to “restrain” excitability within brain circuits (Trevelyan *et al*., 2006; Trevelyan *et al*., 2007; Trevelyan and Schevon, 2013). Ongoing failure of inhibitory restraint also underlies the ability of seizures to perpetuate in time and space. Inhibitory restraint can fail due to reduced GABA release (Zhang *et al*., 2012), GABA_A_R internalization (Goodkin *et al*., 2005), a depolarizing shift in pyramidal E_GABA_ (Lillis *et al*., 2012) or GABAergic interneurons entering a state of depolarization block (Ziburkus *et al*., 2006; Cammarota *et al*., 2013). Knowing which of these mechanisms are involved in SE, and to what extent, will be important for designing optimum strategies for aborting seizures, especially given recent interest in using optogenetic strategies to enhance the action of GABAergic interneurons (Krook-Magnuson *et al*., 2013; Krook-Magnuson and Soltesz, 2015). Using dual whole-cell patch-clamp recordings, we found a very high correlation between hippocampal GABAergic interneuronal and pyramidal cell activity during the LRD phase. Optogenetic activation of the pan-interneuronal population using ChR2 expression driven by the GAD2 promoter revealed that activation of GABAergic interneurons has a broadly similar effect on epileptiform activity as diazepam in our model. Interneurons are inhibitory prior to SLEs and excitatory during SE-like activity. We found that optical activation of the pan-interneuronal population was sufficient to entrain the frequency of epileptiform discharges to the frequency of light activation during the SE-like phase, and that this entrainment is dependent on intact synaptic transmission via GABA_A_Rs. This provides strong evidence about three key aspects of interneuronal function during the early stages of SE: that GABAergic interneurons are able to release GABA, they are not in depolarizing block, and sufficient postsynaptic GABA_A_Rs are present to mediate GABAergic synaptic transmission. However, due to the transient and widespread collapse of the postsynaptic Cl^-^ gradient in pyramidal neurons, GABAergic interneurons are ineffective at curtailing epileptiform discharges, and in fact drive the generation of these events during SE-like activity. Our data complement recent *in vitro* and *in vivo* results, which suggest that various GABAergic interneuronal subtypes may promote the extension of seizures when activated once epileptiform activity has become established (Ellender *et al*., 2014; Sato *et al*., 2017; Magloire *et al*., 2019).

Our observation of excitatory GABA_A_R mediated signalling during the LRD phase explains the loss of inhibitory efficacy and the pro-epileptiform effects of diazepam and the low concentration of phenobarbital we observed during the LRD phase. This supports prior work in dissociated cell cultures which demonstrated that activity-driven changes in the Cl^-^ gradient reduce the inhibitory efficacy of diazepam (Deeb *et al*., 2013). Together, this suggests that benzodiazepines and low concentrations of barbiturates can lose their efficacy in SE even with GABA_A_Rs intact. Interestingly we observed that at high concentrations phenobarbital is able to attenuate and abort SE-like activity. This is likely due to the additional antagonistic effect of high concentrations of phenobarbital on glutamatergic responses via blockade of AMPA and kainate receptors (Macdonald and Barker, 1978; Ko *et al*., 1997; Meldrum and Rogawski, 2007; Nardou *et al*., 2011).

Given our experimental findings, it is worth considering why benzodiazepines are often effective in terminating SE in patients (in our study 52% of patients with SE responded to benzodiazepine administration). Brain slices represent a relatively small, well connected brain circuit, where most areas of the slice are typically involved in SE-like activity (Ellender *et al*., 2014). This means that most pyramidal cells in the slice may experience an activity-dependent collapse of the Cl^-^ gradient, which will render benzodiazepines ineffective during prolonged seizures. In patients with SE, the situation is likely to be different. The proportion of brain circuits affected by activity-dependent Cl^-^ accumulation will vary considerably between patients depending on the seizure type as well as the extent to which different brain areas are recruited into the seizure process. Interactions between areas may also be critical in providing pacemaker drives (Codadu *et al*., 2018), thereby maintaining a high rate of discharge, but breaking this loop by slightly reducing the intrinsic rate at a particular site may be all that is required. In regions where intracellular Cl^-^ and E_GABA_ are still low, benzodiazepines will enhance inhibitory restraint. Therefore, determining the potential cumulative effect of benzodiazepines may involve a dynamic contest between areas where it enhances inhibition and those areas where it is ineffective or promotes hyperexcitability.

Given the demonstrated potential for benzodiazepines to be ineffective, or even promote seizure prolongation in SE, we recommend investigation of alternative or adjunctive therapeutic strategies for terminating SE. These could involve strategies for enhancing Cl^-^ extrusion capacity to help maintain E_GABA_ (Gagnon *et al*., 2013; Alfonsa *et al*., 2016; Moore *et al*., 2017; Magloire *et al*., 2019), or targeting of other brain inhibitory systems including modulating pH (Tolner *et al*., 2011), postsynaptic K^+^ conductance (Zhang *et al*., 2017) or glutamatergic transmission. Our work supports the use of agents which target multiple receptor systems for aborting seizure activity, such as the barbiturate phenobarbital. Given that total duration in status is correlated with morbidity, and that phenobarbital at sufficiently high dosage is highly effective at aborting status epilepticus (Burman *et al*., 2019), we suggest that phenobarbital be re-evaluated as a first-line agent in SE. This is particularly the case as long-term cognitive side effects of once-off phenobarbital use have not been demonstrated.

In summary, our findings support the idea that dynamic network changes and ionic mechanisms likely contribute to the development of persistent seizure activity and that an enhanced understanding of these alterations will guide the development of more effective strategies for treating SE.

## Acknowledgements

We thank Hayley Tomes and Buchule Mbobo who provided assistance with the data collection. We also thank Dr Sharika Raga who assisted in acquiring the phenobarbital used in this study.

---

**Supplementary Figure 1:**
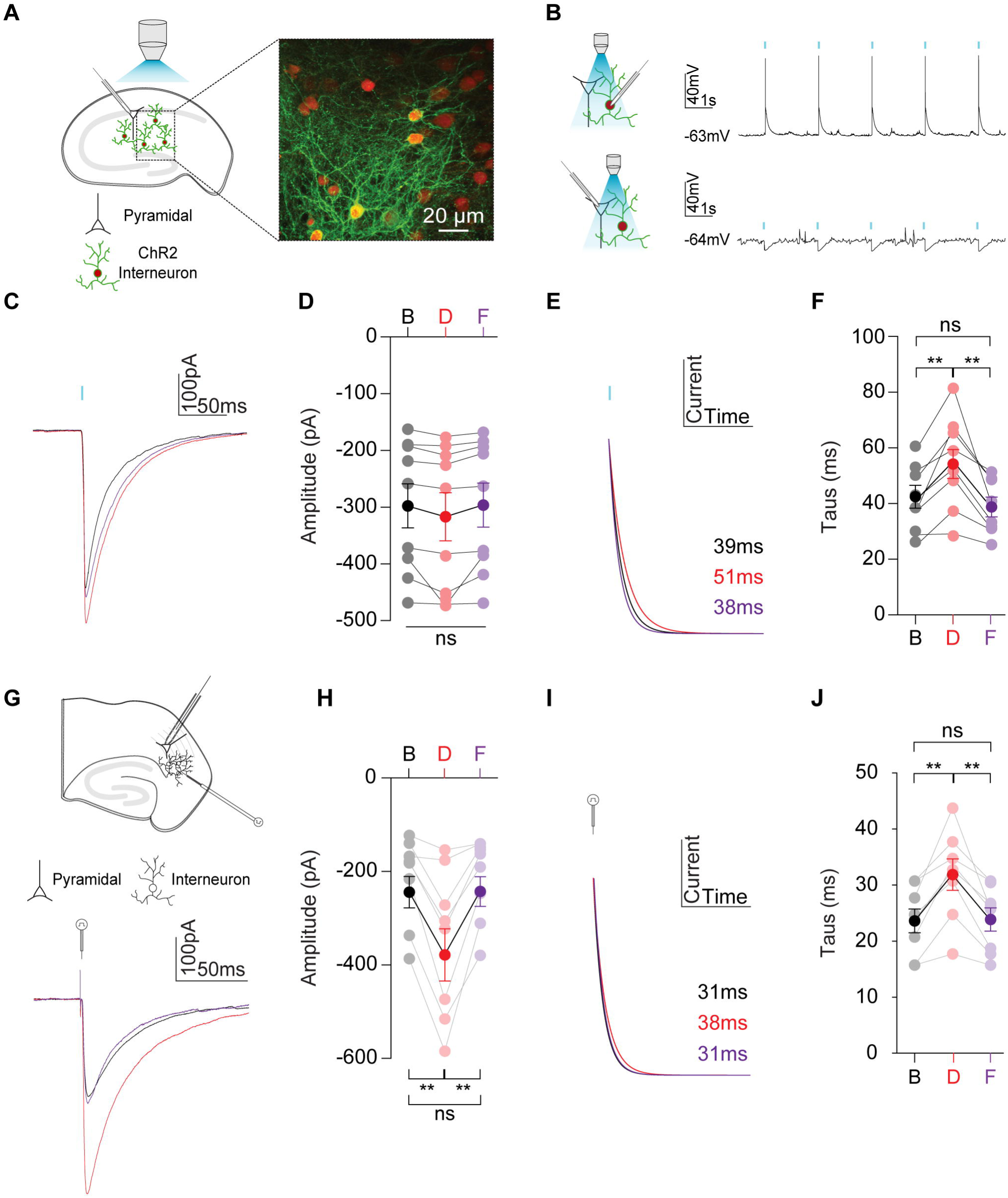
Diazepam enhances GABA_A_R mediated currents in mouse hippocampal organotypic brain slice cultures and acute temporal cortex slices. **A**, Diagram of the experimental setup, involving whole-cell patch-clamp recordings in organotypic slices where Chr2-YFP and tdTomato were expressed in GABA producing (glutamic acid decarboxylase type 2 / GAD2 cells) using the cre-lox system. Inset, confocal microscopy image demonstrating cytoplasmic expression of tdTomato and membrane localisation of ChR2-YFP. **B**, ChR2 activation using a blue LED (470 nm at 17.1 mW, 100ms) reliably elicited action potentials in fluorescent neurons (top) and hyperpolarizing IPSPs in CA1 pyramidal cells (bottom). **C**, Whole-cell voltage clamp recordings with a high Cl^-^ internal (141 mM) containing QX314 (5 μM) and held at −40 mV in the presence of the GABA_B_ receptors blocker, CGP55845 (10 μM), to elicit GABA_A_R synaptic currents (GSCs) following blue light activation of the ChR2-expressing GABAergic interneurons. **D**, The mean amplitude of GSCs calculated from 30 traces from each cell (*n* = 9) recorded under three conditions: baseline (B, black), following 5 min wash-in of 3μM diazepam (D, red), and a further 5 min wash in of the benzodiazepine antagonist 0.4 μM flumazenil (F, purple). We found no significant difference in mean amplitude between the three treatment groups. **E**, The decay phase of GSCs recorded in ‘C’ and normalized to the maximum GSC amplitude. **F**, Population data demonstrating that the decay time constant (tau) was significantly increased in the presence of diazepam (pre-diazepam: mean 42.6 ± *SEM* 4.1 ms vs post-diazepam: mean 56.6 ± *SEM* 5.9 ms, *p* = 0.004, *paired t-test)*. This effect of diazepam was reversed when flumazenil was washed in (post-diazepam: mean 56.6 ± *SEM* 5.9 ms vs flumazenil: 40.3 ± *SEM* 3.6 ms, *p* = 0.004, *paired t-test*) **G**, Horizontal slices (400 μm thick) of mouse temporal lobe including the entorhinal cortex and hippocampal formation were cut using a Compresstome VF200 using a cutting solution composed of (in mM): NaCl (60); KCl (3); NaH2PO4 (1.2); NaHCO3 (23); D-glucose (11); MgCl2 (3); CaCl2 (1) and sucrose (120). Top, diagram of the experimental setup demonstrating whole-cell patch-clamp recordings from layer V pyramidal neurons in entorhinal cortex concurrent with electrical stimulation of afferent fibres using a bipolar stimulation electrode (3ms stim) in the presence of the glutamate receptor blocker, kynurenic acid (2 μM) to isolate GSCs. Bottom, the mean GSC for each cell (*n* = 8) was calculated from 30 traces in each of the different treatment groups: baseline (B, black), diazepam – 3 μM (D, red) and flumazenil - 0.4 μM (F, purple). **H**, Population data of mean GSC amplitudes in each of the treatment groups showing a significant increase in GSC size in the presence of diazepam (pre-diazepam: mean 216.5 ± *SEM* 33.6 pA vs post-diazepam: mean 350.8 ± *SEM* 56.0 pA, *p* = 0.003, *paired t-test)* that was reversed by flumazenil (mean 350.8 ± *SEM* 56.0 pA vs flumazenil: 215.4 ± *SEM* 31.6 pA, *p* = 0.003, *paired t-test*). **I**, Decay phase of GSCs recorded in ‘**G**’ normalized to the maximum GSC amplitude. Deactivation kinetics were fitted by a single exponential function to calculate the time constant (tau) for each condition. **J**, population data of taus for each group showing diazepam significantly increased taus (pre-diazepam: mean 24.0 ± *SEM* 2.1 ms vs post-diazepam: mean 32.3 ± *SEM* 2.8 ms, *p* = 0.001, *paired t-test)* with this effect being reversed by the application of flumazenil (post-diazepam: mean 32.3 ± *SEM* 2.8 ms vs flumazenil: 24.2 ± *SEM* 2.1 pA, *p* = 0.003, *paired t-test*). ns = not significant (p > 0.05); **p <0.01; error bars indicate mean ± SEM.

**Supplementary Figure 2:**
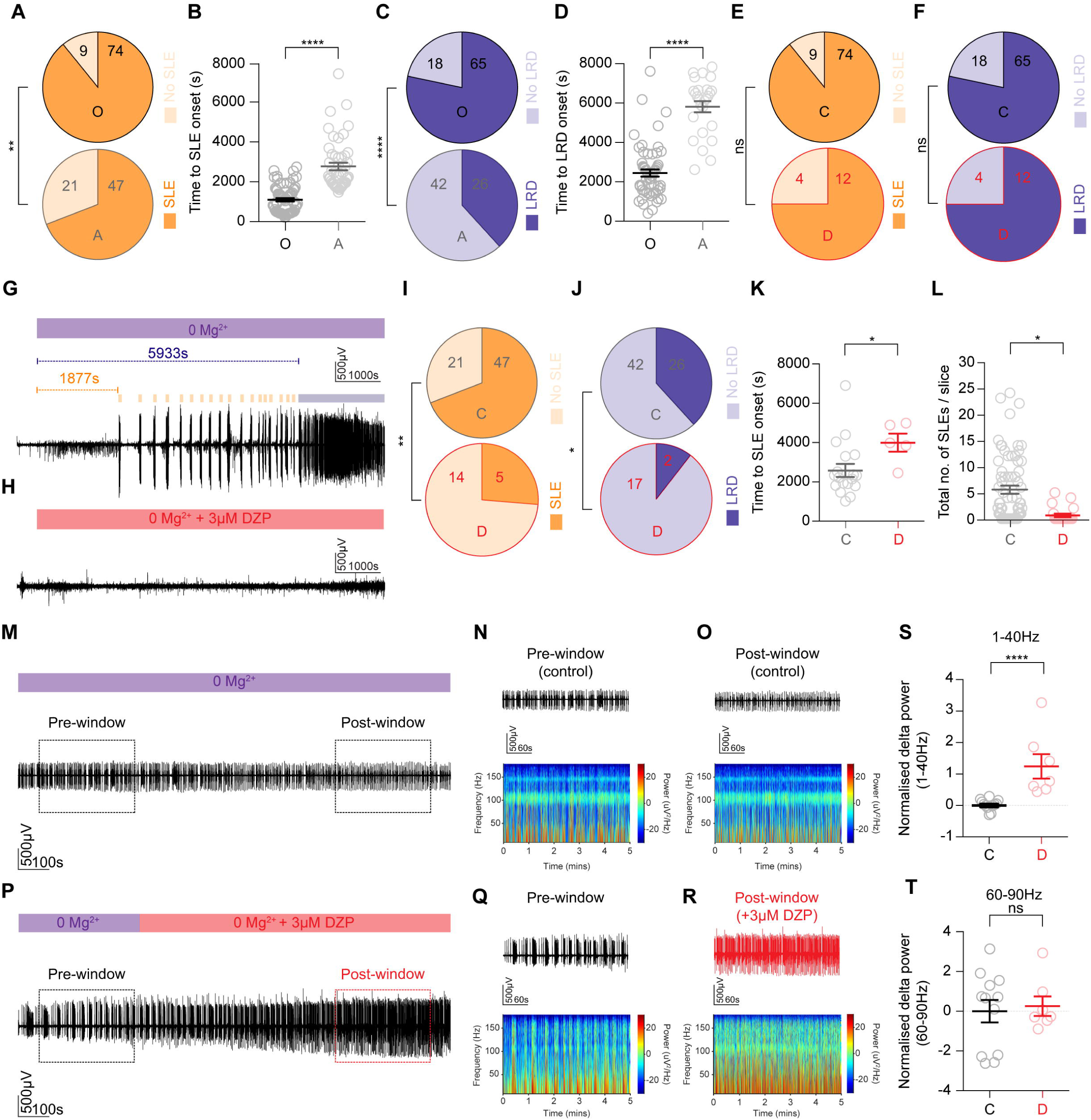
Early application of diazepam has an antiseizure effect while late application is ineffective and augments epileptiform bursting activity in an acute brain slice model of status epilepticus. **A**, Pie charts demonstrating that a larger fraction of organotypic slices generated SLEs following Mg^2+^ withdrawal (top, O) as compared to acute slices (bottom, A) (*p* = 0.004, *Fisher’s exact test*). **B**, Time to 1^st^ SLE was shorter in organotypic as compared to acute slices (organotypic: median 1080 IQR 446.0 – 1573s vs acute: median 2288 IQR 1864 – 3273 s, *p* <0.0001, *Mann-Whitney U-test*). **C**, A greater fraction of organotypic slices successfully entered the LRD phase (top, O) as compared to acute slices (bottom, A) (*p* < 0.0001, *Fisher’s exact test*). **D**, Time to LRD onset was shorter in organotypic as compared to acute slices (organotypic: median 2296 IQR 1506 – 2887 s vs acute: median 6242 IQR 4724 – 6622 s, *p* <0.0001, *Mann-Whitney U-test*). Pie charts demonstrate that in organotypic brain slices, following early wash in of diazepam, a comparable fraction of slices generated SLEs (**E**) and entered LRD (**F**) as compared to controls although diazepam does increase the time to 1^st^ SLE and time to LRD in these slices (see Fig. 2). **G**, Entorhinal cortex LFP recording from a control experiment in acute brain slices where 0 Mg^2+^ was washed in after a 5-10 min baseline period in standard aCSF. Time to SLE onset (orange) and entry into the LRD phase (blue) are shown. **H**, Example recording where early application of diazepam (3 μM) introduced together with 0 Mg^2+^ prevented both SLE generation and entry into LRD in this slice. Pie charts demonstrate that early diazepam application in acute slices as in ‘**H**’, resulted in a smaller fraction of slices generating SLEs (**I**, *p* = 0.001, *Fisher’s exact test*) and entering LRD (**J**, *p* = 0.02, *Fisher’s exact test*) as compared to controls. In acute slices, wash in of diazepam concurrent with 0 Mg^2+^ aCSF extended the time to 1^st^ SLE (**K**, control, *n* = 18: median 2237 *IQR* 1778 – 3287 s vs diazepam, *n* = 5: 3943 *IQR* 3109 – 4903 s, *p* = 0.02, *Mann-Whitney test*) and reduced the total number of SLEs recorded per slice when compared to control slices (**L**, control, *n* = 68: median 3.0 *IQR* 0.0 – 10.5 vs diazepam, *n* = 19: 0.0 *IQR* 0.0 – 5.0, *p* = 0.0001, *Mann-Whitney test*). **M**, Control LFP recording of the LRD phase from an acute brain slice with the 5 min ‘Pre-window’ and ‘Post-window’ used for analysis indicated by dashed rectangles. **N**, ‘Pre-window’ LFP trace (top) with its accompanying spectrogram (bottom). **O**, ‘Post-window’ as in ‘**N**’. **P**, Acute slice LFP recording with diazepam application during LRD and accompanying ‘Pre-window’ (**Q**, prior to diazepam) and ‘Post-window’ (**R**, following diazepam) used for analysis. S, Population data from acute slices showing significantly increased normalised change in power of SE-like activity in the 1-40 Hz bandwidth in diazepam as compared to control slices (control, *n* = 12: median 0.01 *IQR* −0.1 – 0.13 vs diazepam, *n* = 7: 0.75 *IQR* 0.5 – 1.8, *p* < 0.0001, *Mann-Whitney test*). This difference was not observable in the 60-90 Hz frequency range (**T**, control: mean 0.7 *IQR* −2.3 – 1.5 vs diazepam: −0.5 *IQR* −0.5 – 0.9, *p* = 0.8, *Mann-Whitney test*). \**p* ≤ 0.05; \*\**p* ≤ 0.01; ‘ns’, not significant (*p* ≥ 0.05); error bars indicate mean ± SEM.

**Supplementary Figure 3:**
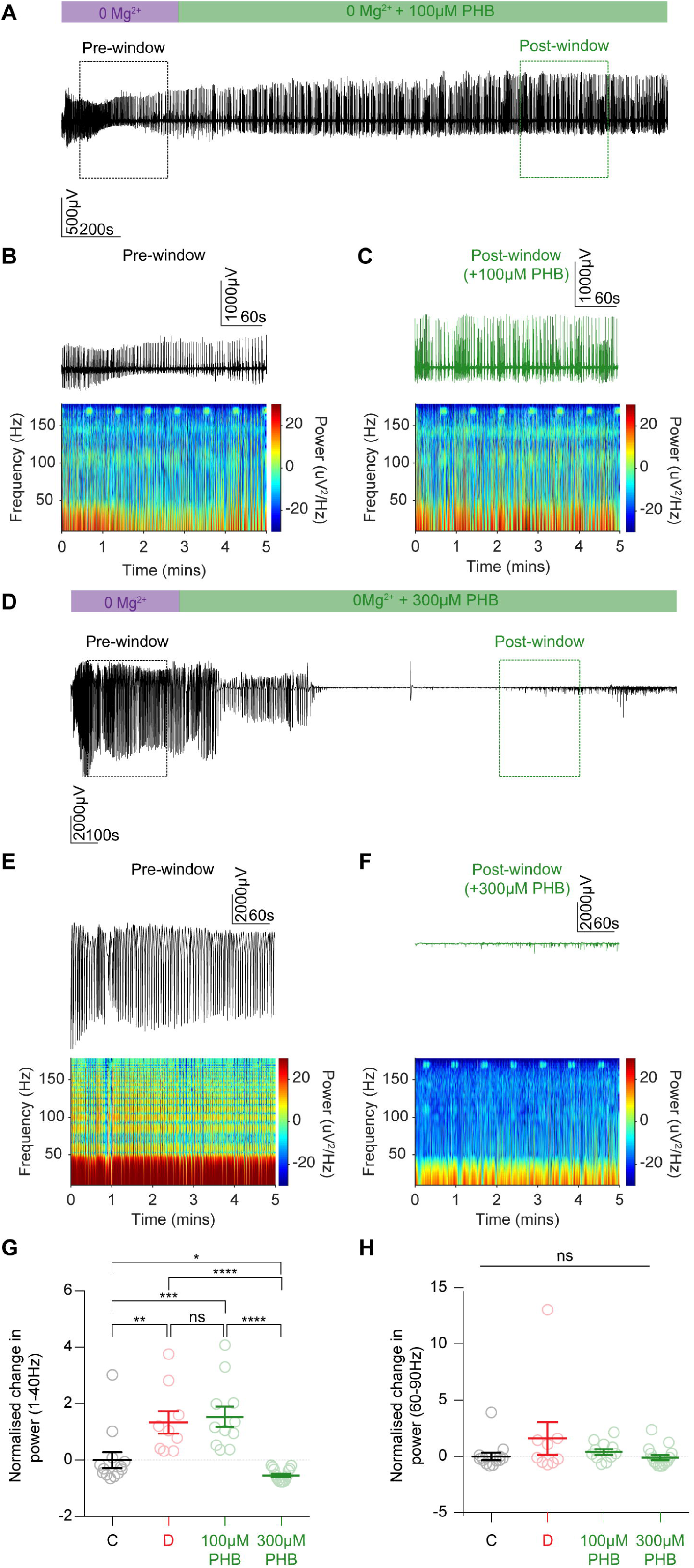
Low concentration phenobarbital exacerbates status epilepticus-like activity whilst a high concentration is antiseizure. **A**, LFP recording form the CA1 area of a hippocampal organotypic brain slice. SE-like activity was elicited following an extended period of Mg^2+^ withdrawal. Dashed rectangles indicate a 5 min ‘Pre-window’ and a 5 min ‘Post-window’ taken after application of 100 μM phenobarbital which were used for power spectral density analysis. **B**, ‘Pre-window’ LFP trace (top) with its accompanying spectrogram (bottom). **C**, ‘Post-window’ as in ‘**B**’. **D, E, F** as in ‘**A**’, ‘**B**’ and ‘**C**’ but with application of a high concentration (300 μM) of phenobarbital during LRD. Note, how in this slice 300 μM phenobarbital abolishes SE-like activity (**F**). **G**, Population data demonstrating that addition of 100 μM phenobarbital during the LRD phase increases the normalised change in power of epileptiform activity in the 1-40 Hz frequency range in a similar manner to 3 μM diazepam when compared to control slices (control, *n* = 13: median - 0.3 *IQR* −0.5– - 0.01 vs 100 μM phenobarbital, *n* = 11: 1.13 *IQR* 0.5 – 2.0, *p* = 0.0003, *Mann-Whitney test*). In contrast, 300 μM phenobarbital significantly reduces the normalised change in power of epileptiform activity compared to controls, demonstrating an antiseizure effect of this dosage (control, *n* = 13: median −0.3 *IQR* −0.5– - 0.01 vs 300 μM phenobarbital, *n* = 15: - 0.6 *IQR* −0.8 – −0.3, *p* = 0.02, *Mann-Whitney test*). H, No significant differences in normalized change in power in the 60-90 Hz range were observed between any of the groups. \**p* ≤ 0.05; \*\**p* ≤ 0.01; \*\*\**p* ≤ 0.001; \*\*\*\**p* ≤ 0.0001; ‘ns’, not significant (*p* ≥ 0.05); error bars indicate mean ± SEM.

**Supplementary Figure 4:**
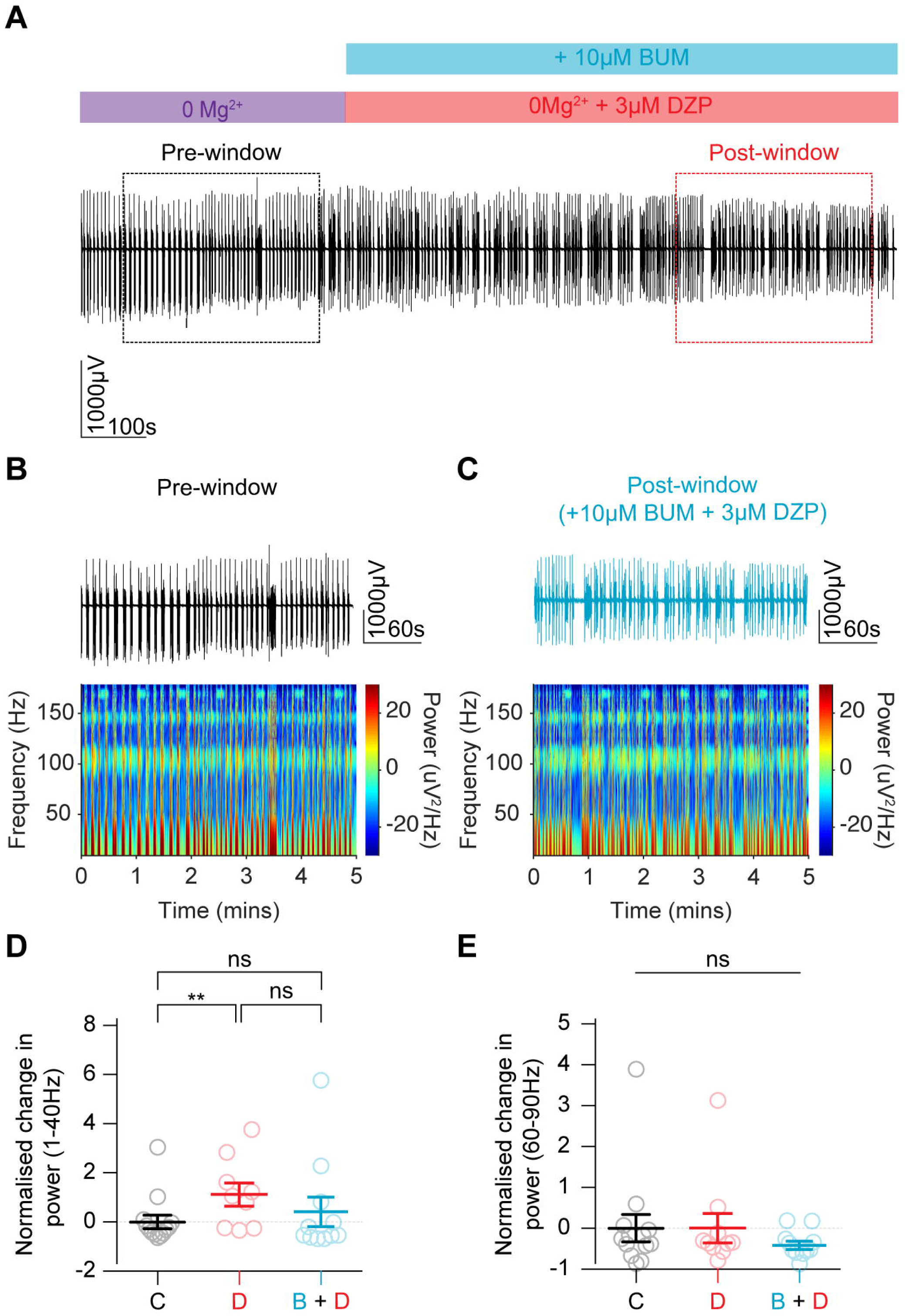
Bumetanide does not rescue the efficacy of diazepam during *in vitro* status epilepticus-like activity. **A**, LFP recording form the CA1 area of a hippocampal organotypic brain slice depicting the LRD phase generated following an extended period of exposure to 0 Mg^2+^ aCSF. Dashed rectangles indicate a 5 min ‘Pre-window’ and a 5 \min ‘Post-window’ taken after application of 10 μM bumetanide in conjunction with 3μM diazepam. **B,** ‘Pre-window’ LFP trace (top) with its accompanying spectrogram (bottom). **C,** ‘Post-window’ as in ‘B’. **D,** Population data showing that addition of 10 μM bumetanide in conjunction with 3 μM diazepam during the LRD phase does not significantly modify the normalised change in power of epileptiform activity in the 1-40 Hz frequency range as compared to diazepam alone or control slices (diazepam only, *n* = 9: median 1.01 *IQR* 0.3 – 2.2 vs diazepam + bumetanide, *n* = 11: median −0.6 *IQR* −0.7 – 0.8, *p* = 0.08, *Mann-Whitney U-test*). **E,** There were no significant differences in normalized change in power in the 60-90 Hz range. \*\**p* ≤ 0.01; ‘ns’, not significant (*p* ≥ 0.05); error bars indicate mean ± SEM.

