## Supplementary Tables for "Excitatory GABAergic signalling is associated with acquired benzodiazepine resistance in status epilepticus"

*Full title*

*Running Title*

GABAergic signalling in status epilepticus

*Keywords*

status epilepticus, chloride, GABAA receptors, inhibition, seizures

*Authors*

Richard J. Burman^1,2,3^, Joshua Selfe^1^, John Hamin Lee^1^, Maurits van den Burg^1^, Alexandru Calin^3^, Neela K. Codadu^4^, Rebecca Wright^3^, Sarah E. Newey^3^, R. Ryley Parrish^4^, Arieh A. Katz^5^, Joanne M. Wilmshurst^2^, Colin J. Akerman^3^, Andrew J. Trevelyan^4^, Joseph V. Raimondo^1^

^1^Division of Cell Biology, Department of Human Biology, Neuroscience Institute and Institute of Infectious Disease and Molecular Medicine, Faculty of Health Sciences, University of Cape Town, Cape Town, South Africa

^2^Department of Paediatric Neurology, Red Cross War Memorial Children’s Hospital, Cape Town South Africa

^3^Department of Pharmacology, University of Oxford, Oxford, United Kingdom

^4^Institute of Neuroscience, Newcastle University, United Kingdom

^5^Division of Medical Biochemistry, Department of Integrated Biomedical Sciences, Infectious Disease and Molecular Medicine, Faculty of Health Sciences, University of Cape Town, Cape Town, South Africa

**Supplementary Tables:**

| **Supplementary Table 1:**  **Cohort demographics and seizure history at first admission** | | | |
| --- | --- | --- | --- |
|  | **bzpS group**  (*n* = 52) | **bzpR group**  (*n* = 49) | ***p*** |
| **Gender** | | | |
| female | 33 (64%) | 25 (51%) | 0.23 |
| male | 19 (36%) | 24(49%) |  |
| **Seizure history** | | | |
| Previous admission for seizures | 29 (56%) | 22 (45%) | 0.68 |
| Known epileptic | 28 (54%) | 19 (39%) | 0.23 |

Significant associations determined by *Fisher’s exact test*. ‘bzpS’, benzodiazepine sensitive; ‘bzpR’, benzodiazepine resistant.

| **Supplementary Table 2:**  **Benzodiazepine sensitivity across multiple admission for CSE** | | | |
| --- | --- | --- | --- |
|  | **bzpS group**  (*n* = 75) | **bzpR group**  (*n* = 69) | ***p*** |
| First admission | 52 (69%) | 49 (71%) | 0.22 |
| Second admission | 18 (24%) | 12 (17%) | 0.20 |
| Third admission | 5 (7%) | 6 (8%) | 1 |
| Fourth admission | 0 | 1 (2%) | 1 |
| Fifth admission | 0 | 1 (2%) | 1 |

Significant associations determined by *Fisher’s exact test*. ‘bzpS’, benzodiazepine sensitive; ‘bzpR’, benzodiazepine resistant; ‘CSE’, convulsive status epilepticus.

| **Supplementary Table 3:**  **Presentation of paediatric convulsive status epilepticus for all admissions** | | | |
| --- | --- | --- | --- |
|  | **bzpS group**  (*n* = 75) | **bzpR group**  (*n* = 69) | ***p*** |
| **Age at admission** – median (IQR) | 38 (18 – 69) | 24 (14 – 64) | 0.08^*^ |
| **Type of SE** | | | |
| Continuous | 37 (49%) | 36 (52%) | 0.74 |
| Intermittent | 38 (51%) | 33 (48%) |  |
| **Semiology** | | | |
| Focal onset evolving into bilateral SE | 15 (20%) | 13 (19%) | 0.19^#^ |
| Generalised | 52 (69%) | 41 (59%) |  |
| Unknown focal or generalised | 8 (11%) | 15 (22%) |  |
| **Febrile CSE** | 12 (16%) | 15 (22%) | 0.59 |

Continuous SE was defined as a convulsive seizure that lasted longer than 5min. Intermittent SE was defined as multiple discrete seizures between which there was no extended period of recovery between events. The classifications of SE type and semiology follow the most recent multiaxial diagnostic criteria for SE (Trinka *et al.*, 2015). The diagnosis of Febrile CSE was only made if it was apparent that the CSE had been provoked by hyperthermia (>38.4 degrees Celsius) with there being no evidence of acute central nervous system diseases or prior history of afebrile seizures (Hesdorffer *et al.*, 2012). Significant associations determined by *Fisher’s exact test*. #Significant difference in proportions determined by Chi-squared test. ^*^denotes where Mann Whitney U test was used to compare nonparametric data. ‘bzpS’, benzodiazepine sensitive; ‘bzpR’, benzodiazepine resistant; ‘CSE’, convulsive status epilepticus.

| **Supplementary Table 4:**  **Contingency table showing benzodiazepine response**  **against duration of CSE** | | | |
| --- | --- | --- | --- |
| Benzodiazepine response | *up to 1 hour* | *>1 hour* | *Total* |
| *Sensitive* | 67 | 8 | 75 |
| *Resistant* | 33 | 36 | 69 |
| *Total* | 100 | 44 | 144 |

‘CSE’, convulsive status epilepticus.
